# An integrin centered complex coordinates ion transport and pH to regulate f-actin organization and cell migration in breast cancer

**DOI:** 10.1101/2024.07.02.601509

**Authors:** Chiara Capitani, Jessica Iorio, Elena Lastraioli, Claudia Duranti, Giacomo Bagni, Ginevra Chioccioli Altadonna, Rossella Colasurdo, Giorgia Scarpellino, Scott P. Fraser, Andrea Becchetti, Mustafa B. A. Djamgoz, Annarosa Arcangeli

## Abstract

Reciprocal signaling between the Tumor Microenvironment (TME) and cancer cells regulates abnormal proliferation, migration and pro-metastatic behavior. Major player in such interaction is integrin-mediated cell adhesion to the Extracellular Matrix (ECM). Integrin receptors organize signaling hubs constituted by multiprotein membrane complexes often comprising ion channels and transporters. We studied whether and how integrin-centered multiprotein complexes control cell behavior in Breast Cancer (BCa) cell populations with different molecular characteristics. BCa cells were cultured onto the ECM protein fibronectin (FN), to trigger β1 integrin activation. Through biochemical, immunofluorescence and electrophysiological experiments we provide evidence of a novel signaling pathway that involves a β1 integrin-centered plasma membrane complex formed by different transport proteins: the hERG1 K^+^ channel, the neonatal form of the Na^+^ channel Na_V_1.5 (nNa_V_1.5) and the Na^+^/H^+^ antiporter NHE1. The NHE1/hERG1/β1/nNa_V_1.5 complex was found on the plasma membrane of BCa cells, and particularly of Triple Negative Breast Cancer (TNBCa). When engaged by cell adhesion to FN, such membrane complex recruited the cytoskeletal actin-binding protein a-actinin1 and stimulated NHE1-mediated cytoplasmic alkalinization. Thus, the multiprotein complex activation affected TNBCa migration and invasiveness by stimulating f-actin organization directly (through α-actinin1) and indirectly (by intracellular alkalinization). The contribution of both hERG1 and nNa_V_1.5 was essential, as the adhesion-dependent signaling pathway and its functional consequences were inhibited by blocking either channel with, respectively, E4031 and TTX, or by applying RNA silencing procedures. The contribution of hERG1 to the structural integrity of the membrane complex appeared to be critical, as the adhesion-dependent signals were hampered by harnessing the hERG1/β1 integrin complex with a single chain bispecific antibody (scDb-hERG1-β1) which disrupts the macromolecular complex without blocking the K^+^ current, as well as by E4031, which impairs the complex formation by blocking the channel in the open state.

In conclusion, we revealed that integrin-centered macromolecular complexes in BCa cells recruit a battery of ion transport proteins that cooperate in modulating different aspects of the downstream signals that lead to malignant behavior. This complex could be targeted to develop novel therapeutic strategies for one of the most difficult-to-treat cancers, i.e. TNBCa.

## INTRODUCTION

Dysregulation of cell migration promotes the development of invasion and metastasis in cancer^1^ and represents one of the hallmarks of cancer^2^. Cell motility is initiated in response to growth factors or chemokines that stimulate the formation of plasma membrane protrusions whose direction is regulated by local Ca^2+^ levels^3,4^. These protrusions, in the form of lamellipodia or filopodia, are driven by the polymerization of f-actin which not only represents a simple regulator of mechanical properties of the cell but also behaves as a platform for signaling pathways^5^. In metastatic cancer cells membrane protrusions are organized in special structures named invadopodia which allow to cross physiological barriers and in turn to trigger the metastatic process^1^. All the above structures that are linked to cell motility and invasion are stabilized by adhesions to Extracellular Matrix (ECM) proteins through specific cell receptors. In these processes, integrins are pivotal since they link the plasma membrane to the f-actin cytoskeleton^6,7^ but also transduce intracellular signaling pathways hence working as bidirectional signal transducers^8^.

Integrin receptors coordinate intracellular signaling through the formation of plasma membrane hubs constituted by multiprotein plasma membrane complexes which often comprise ion channels and transporters^9^. Wide evidence is now available for the *human ether-á-go-go-related gene* (hERG1) K^+^ channel, encoded by the *KCNH2* gene. The complex between the β1 integrin subunit and hERG1 is highly present in cancer cells and operates as a supramolecular signaling hub^10^. The complex modulates cell migration through the re-organization of f-actin in Pancreatic Ductal Adeno Carcinoma (PDAC) and in Colorectal Cancer (CRC) cells^10^. In CRC cells ECM-triggered, integrin-mediated cell migration also depend on intracellular alkalinization through an increased H^+^ efflux via the Na^+^/H^+^ exchanger NHE1^11^.

hERG1 is also expressed in Breast Cancer (BCa) cells^12,13^ and primary samples^14^. In the Triple Negative Breast Cancer (TNBCa) cell line MDA-MB-231 hERG1 is physically associated with the β1 integrin subunit, to form a hERG1/β1 integrin complex which enhances the metastatic potential *in vivo*^15^. BCa cells and primary samples also show an upregulation of the 1.5 subtype of Voltage Dependent Sodium channels (Na_V_1.5) encoded by the *SCN5A* gene. In BCa cells, and in particular in TNBCa cells, Na_V_1.5 promotes invasiveness and metastasis^16,17^ through mechanisms which also involve the Na^+^/H^+^ exchanger NHE1 (reviewed in Luo et al^18^). Interestingly, BCa cells preferentially express the neonatal splice variant of Na_V_1.5, which potentiates BCa cell invasiveness *in vitro*^19^.

The present paper was aimed at studying the hERG1/β1 integrin complex in BCa cells and how it regulates the f-actin cytoskeleton and cell migration. We found that the complex (i) is present in several BCa cell lines, (ii) its formation is triggered by cell adhesion to the ECM protein FN and (iii) recruits both the neonatal form of the Na^+^ channel Na_V_1.5 (nNa_V_1.5) and the Na^+^/H^+^ antiporter^20^. The NHE1/hERG1/β1/nNa_V_1.5 complex is present on the plasma membrane of BCa cells, in particular in TNBCa. The expression of the two ion channels present in the complex is mutually regulated. The complex engagement by cell adhesion to FN stimulates a NHE1-mediated cytoplasmic alkalinization, which modulates f-actin organization. This in turn controls TNBCa migration and invasiveness. All this pathway is impaired by blocking either hERG1 or nNa_V_1.5 which also cause complex disassembly. The same result is obtained by harnessing the hERG1/β1 integrin complex through a single chain bispecific antibody (scDb-hERG1-β1)^21,22^.

## RESULTS

### A hERG1/**β**1 integrin macromolecular complex is present in BCa cells of different molecular subtype

We previously showed that a hERG1/β1 integrin complex is present on the plasma membrane of cancer cells, including MDA-MB-231 BCa cells^15^. We first studied whether this complex was present in different BCa cells, analyzing four human cell lines with different molecular characteristics: MCF-7 (luminal A, weakly metastatic), SKBR3 (HER2 positive, weakly metastatic), MDA-MB-468 and MDA-MB-231 (TNBCa, strongly metastatic)^23^. All the analyses were done on cells seeded for 2 hours on fibronectin (FN) as in Duranti and Iorio et al^10^. We first determined the levels of the *KCNH2* transcript (by RQ-PCR) and of the hERG1 protein by WB and IF. For IF we used a specific antibody recognizing an extracellular epitope of the channel (mAb-hERG1, Table S1)^24,25^. *KCNH2* was expressed in all the cells at varying levels: MDA-MB-468 and MDA-MB-231 cells showed the highest levels, while SKBR3 and MCF-7 cells the lowest (**Figure 1A**). The hERG1 protein was expressed in all the native BCa cell lines at comparable level (**Figure 1B**). When the plasma membrane localization of the channel was analyzed by IF, MDA-MB-468 cells showed the highest levels, MCF-7 the lowest and MDA-MB-231 and SKBR3 intermediate levels (**Figure 1C**). Proteins from all the cell lines were immunoprecipitated with either a monoclonal antibody recognizing extracellular epitopes of the β1 integrin (mAb-TS2/16) (IP: β1 integrin)^26^ or the mAb-hERG1 (IP: hERG1). Polyclonal antibodies directed against intracellular epitopes of the two proteins were used to label the WBs (**Figure 1D**). Co-immunoprecipitation (co-IP) data were quantified by densitometry as detailed in Materials and Methods (see the bar graph showing the densitometric data on the right of Figure 1D). Overall, a hERG1/β1 integrin complex occurs in BCa cells once seeded on FN for 2 hours, especially in TNBCa cells. As previously shown in PDAC and CRC cells^10^, also in BCa cells the complex was sensitive to (meaning was dissociated by) treatment with either the classical hERG1 blocker E4031 which impairs the complex formation by blocking the channel in the open state^10,15^ or the bispecific antibody which specifically targets the hERG1-β1 integrin complex (scDb-hERG1-β1) and disrupts the macromolecular complex without blocking the current^10,22^. This was evident from either co-IP (**Figure 1E**) or IF experiments, where the fluorescence intensity obtained after labeling the complex with scDb-hERG1-β1 was determined (**Figure 1F**). Both E4031 and scDb-hERG1-β1 treatments reduced the complex formation, although the effect of scDb-hERG1-β1 was stronger. This occurred only on cells seeded on FN, which triggered the activation of the β1 integrins, but not on cells seeded onto an integrin-independent substrate, BSA. Data relative to TNBCa cells MDA-MB-231 and MDA-MB-468 are in **Figure 1E and 1F**, data relative to MCF-7 and SKBR3 cells are shown in **Supplementary S1B**).

**Figure 1.**
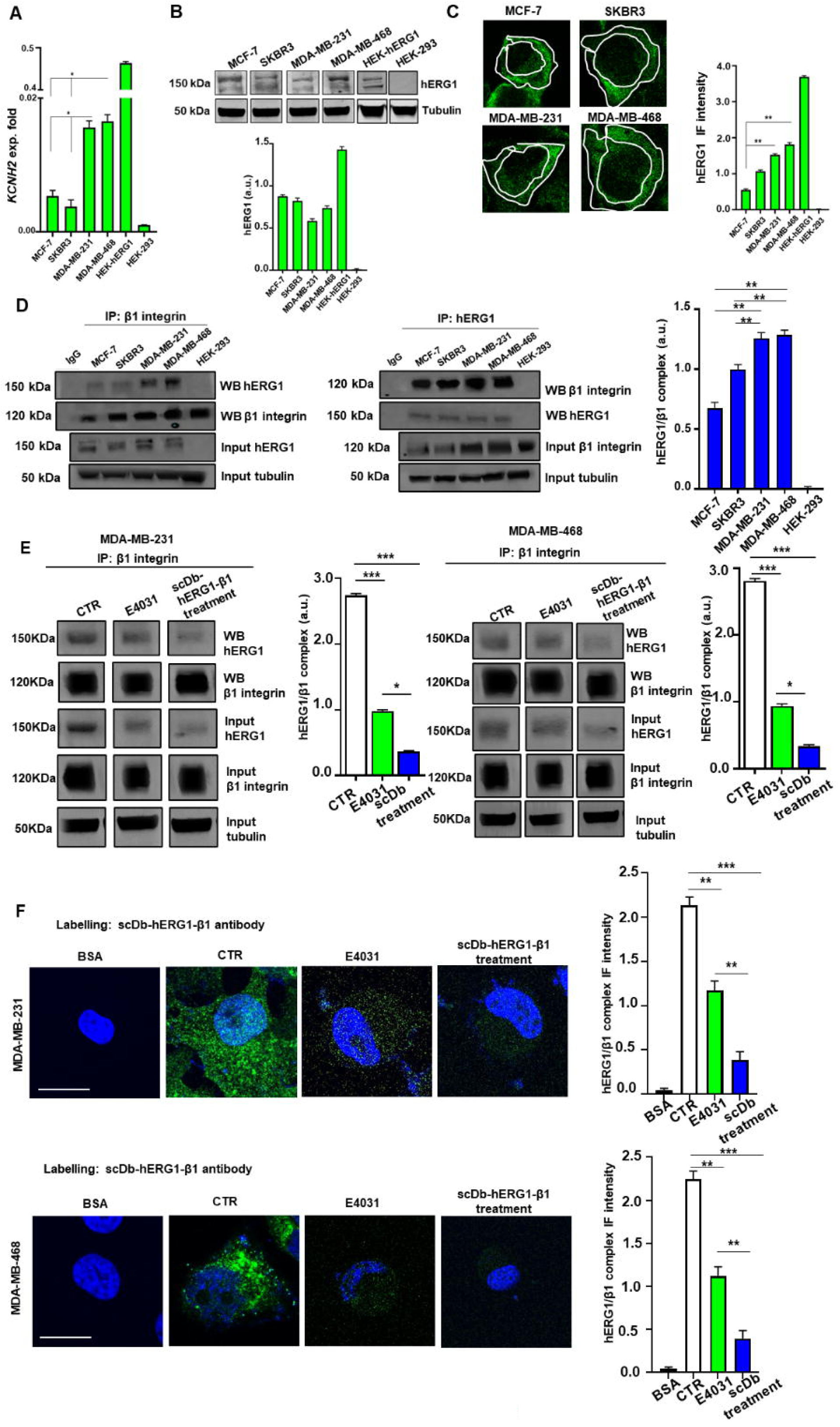
The hERG1/ β1 integrin complex is expressed in breast cancer cell lines of different molecular subtype. **A)** Expression levels of the *KCNH2* transcripts, determined by Real Time PCR in different breast cancer cell lines (MCF-7, SKBR3, MDA-MB-231, MDA-MB-468), HEK-hERG1, and HEK-293 cells. HEK-hERG1 were used as positive control for *KCNH2*, HEK-293 as negative control. Cells were seeded onto Fibronectin (FN) for 2 hours. Data, reported as 2-DCt, are mean values ± s.e.m. (n=3). **B**) Expression levels of the hERG1 protein determined by Western Blot (WB) in breast cancer cell lines (MCF-7, SKBR3, MDA-MB-231, MDA-MB-468), HEK-hERG1 and HEK-293 cells. HEK-hERG1 were used as positive control for hERG1, HEK-293 as negative control for the two proteins. Cells were seeded onto FN for 2 hours. Representative WB (left panel) and densitometric analysis (right panel) are reported. Data are mean values ± s.e.m obtained in three independent experiments. a.u.= arbitrary units. **C)** Membrane expression levels of hERG1, determined by Immunofluorescence (IF) in breast cancer cell lines (MCF-7, SKBR3, MDA-MB-231, MDA-MB-468). Cells were seeded onto FN for 2 hours. Cells were stained for hERG1 (in green) with mAb-hERG1 primary antibody conjugated with AlexaFluor-488. Representative images (scale bar 100 µm) and quantification are reported. The quantification was performed considering only the membrane signal, highlighted by the white masks shown in the pictures. For each condition the fluorescence relative to 20 cells was analyzed. For total IF signal, we measured the fluorescence intensity for each cell, dividing it by the cell area, to obtain a final mean fluorescence (± standard error mean). Full images are reported in **Supplementary Figure S1A**. Bar graph of normalized WB optical densities to the IF cytoplasmic values of hERG1. a.u.= arbitrary units. Pictures were taken on a confocal microscope (Nikon TE2000, Nikon; Minato, Tokyo, Japan). ImageJ software was used to analyse the images. **D)** Co-immunoprecipitation (co-IP) between hERG1 and β1 integrin in BCa cells (MCF-7, SKBR3, MDA-MB-231, MDA-MB-468) and HEK-293 (negative control) seeded onto FN for 2 hours. Immunoprecipitation was performed with a monoclonal antibody recognizing extracellular epitopes of the β1 integrin subunit (TS2/16) and a monoclonal antibody recognizing hERG1 (mAb-hERG1), left panels. Immunoprecipitation was performed with monoclonal antibody recognizing extracellular epitopes of hERG1, (right panel). Left panels: representative experiment; right panels: densitometric analysis (n=3). Data are reported as a.u. (arbitrary unit). **E)** Co-immunoprecipitation of hERG1 with β1 integrins at T_120_. Immunoprecipitation was performed with a monoclonal antibody recognizing extracellular epitopes of the β1 integrin subunit (TS2/16). Representative blots are in the left panels, the densitometric data are in the right panels. Data are mean values ± s.e.m. F) hERG1/β1 complex formation evaluated by staining with fluorescent scDb–hERG1–β1 in MDA-MB-231 and MDA-MB-468 cells seeded on FN and BSA as negative control. Representative IF images at T_120_ (left panel) (scale bars: 100 μm) and IF densitometric analysis (right panel) are reported. Full images are reported in **Supplementary Figure S1B**. Pictures were taken on a confocal microscope (Nikon TE2000, Nikon; Minato, Tokyo, Japan). ImageJ software was used to analyse the images. *P < 0.05 and **P < 0.01 and *** P < 0.001. IF = immunofluorescence. IP = immunoprecipitation.

We then determined the localization of the hERG1/β1 integrin complex in BCa cells. We first focused on focal adhesions (FAs), i.e. pivotal localization of integrins^27^, testing whether the hERG1/β1 integrin complex co-immunoprecipitated/co-localized with paxillin^28^. Indeed, co-IPs revealed the presence of paxillin in the β1 integrin IPs containing hERG1 (**Figure 2A**). Co-localization of hERG1 and paxillin, evidenced by IF experiments, was quantified as the Manders’ Overlapping Coefficient (MOC). Indeed hERG1 (**Figure 2B**, upper panels, and related MOC values on the bottom of each picture) as well as the complex, labelled with a bispecific antibody recognizing the hERG1/β1 integrin complex, scDb-hERG1-β1, Table S1^10^ (**Figure 2B**, lower panels, and related MOC values on the bottom of each picture.) with paxillin. We then studied whether the hERG1/β1 integrin complex in FAs recruited cytoskeletal proteins such as vinculin, α-actinin 1 or f-actin ^29,30^. Neither f-actin nor vinculin were present in the co-IP of hERG1 and β1 integrin, while α-actinin1 co-immunoprecipitated with the β1 integrin and hERG1 in all the four cell lines, in particular in TNBCa cells (**Figure 2A**). Silencing hERG1 and immunoprecipitating the proteins with mAb-TS2/16 did not affect the co-IP with α-actinin (**Figure 2C**), whereas silencing β1 and immunoprecipitating with the mAb-hERG1 resulted in the withdrawal of both α-actinin1 and hERG1 from the complex (**Figure 2D**).

**Figure 2.**
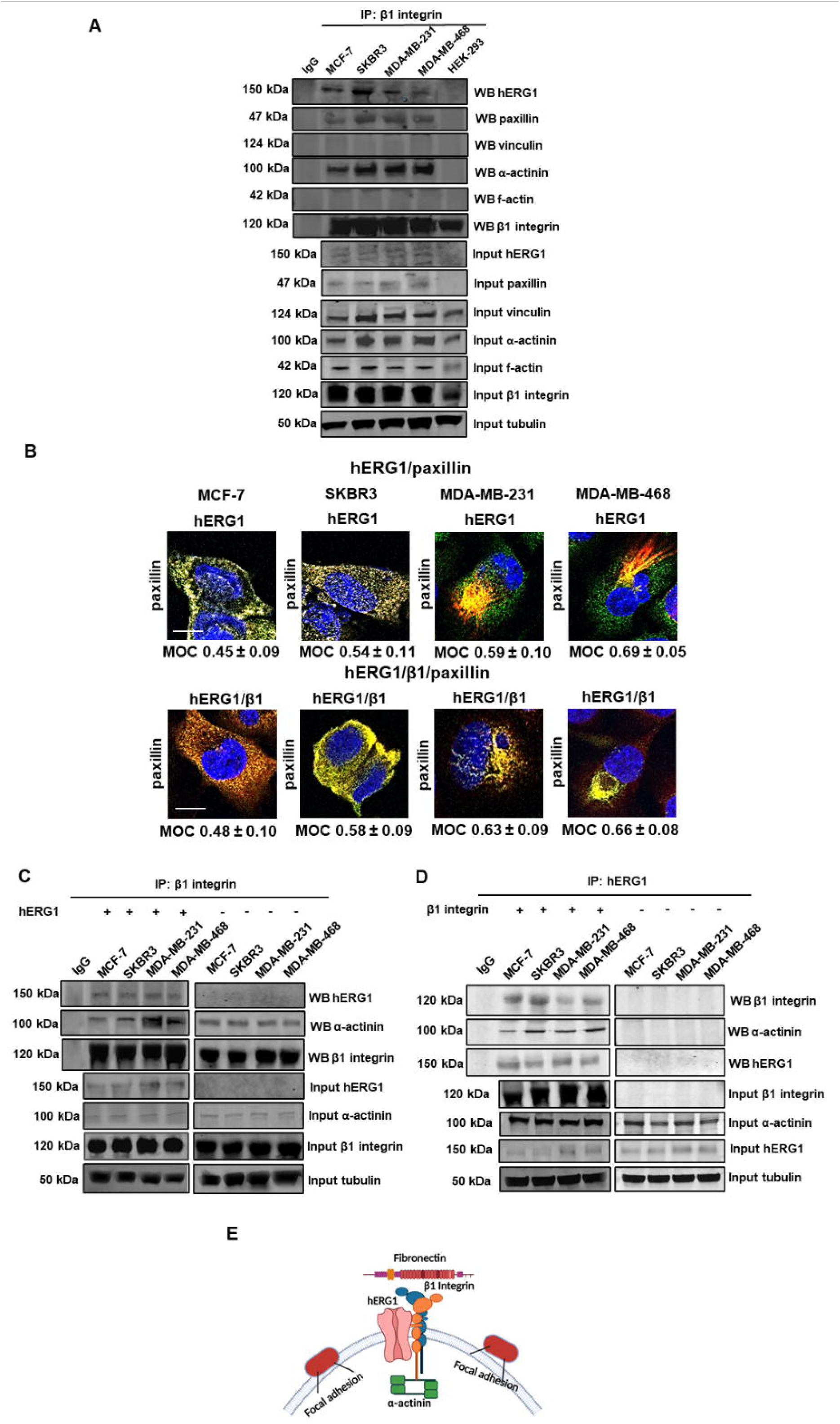
The hERG1/β1 integrin complex is expressed in BCa cells, localizes in FAs and recruits α-actinin1 directly linked to β1 integrin. **A)** Co-IP between β1 integrin and vinculin, paxillin, α-actinin, f-actin in BCa cells (MCF-7, SKBR3, MDA-MB-231, MDA-MB-468) and HEK-293 (negative control) seeded onto FN for 2 hours. Immunoprecipitation was performed with monoclonal antibody recognizing extracellular epitopes of the β1 integrin subunit (TS2/16). Data are reported as a. u. (arbitrary unit). **B)** Representative IF images (scale bar: 100 μm) of the co-localization of hERG1 (in green probed with mAb-hERG1 conjugated with AlexaFluor-488) and paxillin (in red probed with mAb-paxillin and secondary Ab conjugated with AlexaFluor-546) (top panel), and hERG1/β1 (in green probed with scDb-hERG1/β1 conjugated with AlexaFluor-488) and paxillin (in red probed with mAb-paxillin and secondary Ab conjugated with AlexaFluor-546) (bottom panel) in breast cancer cell lines (MCF-7, SKBR3, MDA-MB-231, MDA-MB-468) seeded onto FN for 2 hours. MOC reporting hERG1/paxillin and hERG1/β1/paxillin correlations is showed below each image. For each condition the fluorescence relative to 20 cells was analyzed. For total IF signal, we measured the fluorescence intensity for each cell, dividing it by the cell area, to obtain a final mean fluorescence (± standard error mean). Full images are reported in **Supplementary Figure S2A**. Pictures were taken on a confocal microscope (Nikon TE2000, Nikon; Minato, Tokyo, Japan). ImageJ software was used to analyse the images. **C)** Co-IP between hERG1, α-actinin1, and β1 integrin in BCa cells transiently silenced for hERG1. **D)** Co-IP between hERG1, α-actinin1, and β1 integrin in BCa cells transiently silenced for β1 integrin. Both in C) and D) BCa cells (MCF-7, SKBR3, MDA-MB-231, MDA-MB-468) were seeded onto FN for 2 hours. Immunoprecipitation was performed with monoclonal antibody TS2/16 (C) and monoclonal antibody for hERG1 (mAb-hERG1) (D). Data are mean values ± s.e.m obtained in three independent experiments. a.u.= arbitrary units. **E)** Model representation of the macromolecular complex comprising hERG1, β1 integrin and α-actinin1. Created with BioRender.com. MOC = Manders’ Overlapping Coefficient. IP = immunoprecipitation. FN = fibronectin.

In conclusion, the hERG1/β1 integrin complex is present in BCa cells, especially in TNBCa, localizes in FAs with paxillin and recruits α-actinin1, which is linked to the β1 integrin (**Figure 2E**).

### The hERG1/**β**1 integrin macromolecular complex recruits nNa_V_ channels in BCa cells: the hERG1/**β**1/nNav1.5 complex

Given the high expression and relevant role of Na_V_ channels in BCa cells^17,31^, we analyzed weather the hERG1/β1 integrin complex recruited the Na_V_1.5 channel protein in BCa cells. We followed the same procedure as above, first determining the levels of the *SCN5A* transcript (by RQ-PCR) and of the Na_V_1.5 protein by WB and IF. In particular we quantified the expression of either the adult or the neonatal form of the channel, *aSCN5A* and *nSCN5A*, the latter been expressed in BCa cells (Brackenbury et al, 2007). While *aSCN5A* was hardly detectable, the expression of *nSCN5A* was higher amongst the native cells. MDA-MB-468 and SKBR3 showed a higher expression level compared to MDA-MB-231 and MCF-7 cells (**Figure 3A**). The expression of the two transcripts in HEK cells stably transfected with either the two *SCNA5* cDNAs (EBNA-aNa_V_1.5 and EBNA-nNa_V_1.5 cells, respectively) are also shown in **Figure 3A**. Since *nSCN5A* was dominant we investigated only this isoform in the following experiments. The expression levels of nNa_V_1.5 protein were determined by WB and IF, using a newly developed antibody specific for nNa_V_1.5: mAb-nNa_V_1.5 which recognizes an extracellular epitope of the channel (Duranti et al, personal communication, January 2025). The WB data showed the presence of two bands: a band weighing 227 kDa and a band of higher molecular weight (∼245 kDa). According to what described for the adult form of Na_V_1.5^32^ the two bands correspond to the partially or un-glycosylated, immature form and to the fully glycosylated, mature form of the protein, respectively. All BCa cell lines expressed the protein in comparable levels when quantified (**Figure 3B**). The presence and plasma membrane localization of the channel was confirmed by IF. With this technique MDA-MB-231 and MDA-MB-468 cells showed the highest levels, MCF-7 the lowest (**Figure 3C** and the bar graph on the right).

**Figure 3.**
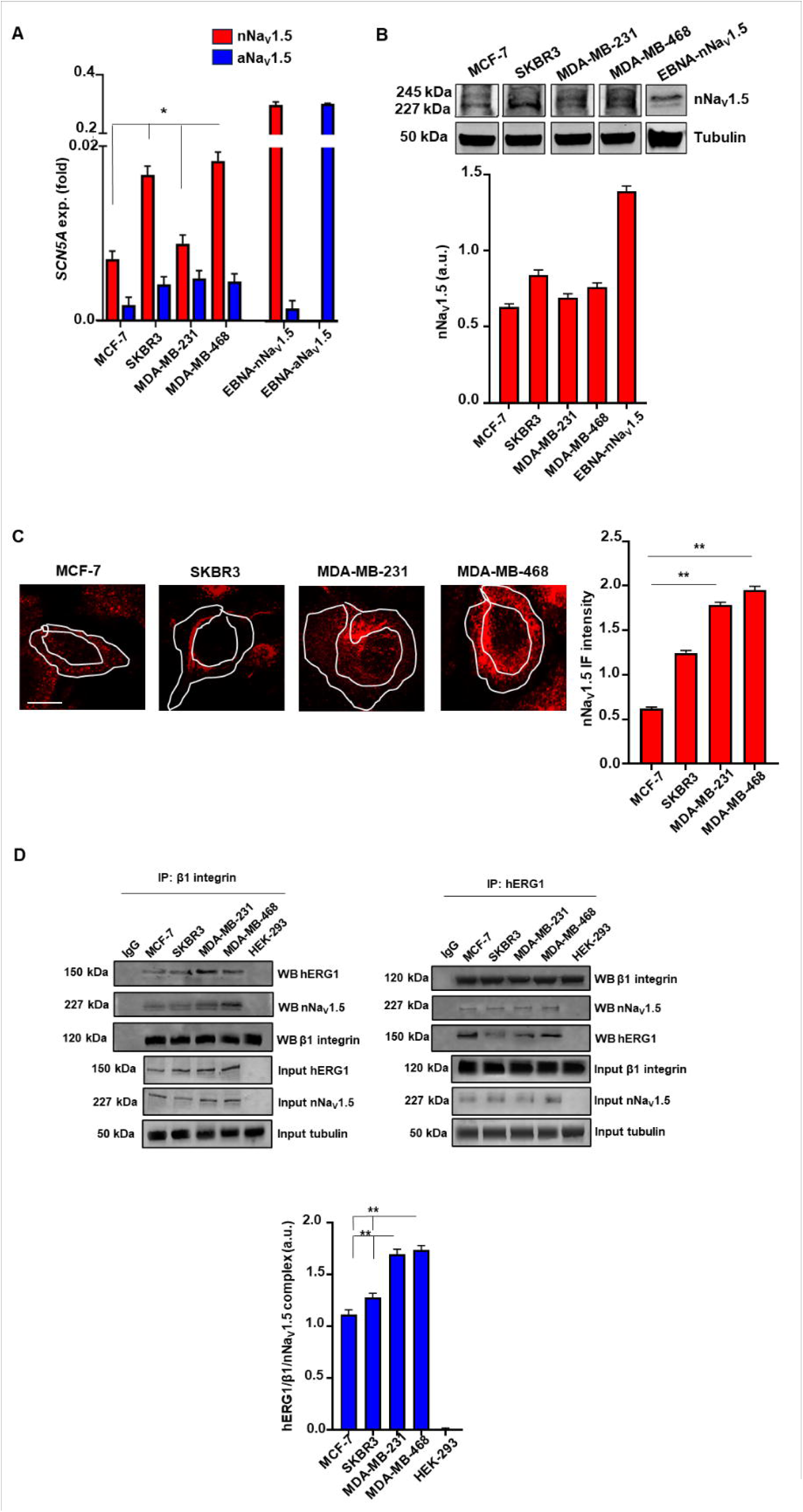
nNa_V_1.5 is expressed in breast cancer cell lines of different molecular subtypes, in focal adhesions and is co-expressed with the hERG1/β1 integrin complex. **A)** Expression levels of the *nSCN5A* and *aSCN5A* transcripts determined by Real Time PCR in different breast cancer cell lines (MCF-7, SKBR3, MDA-MB-231, MDA-MB-468), EBNA-nNa_V_1.5 and EBNA-aNa_V_1.5. EBNA-nNa_V_1.5 were used as positive control for *nSCN5A* while EBNA-aNa_V_1.5 were used as negative control. Cells were seeded onto Fibronectin (FN) for 2 hours. Data, reported as 2-DCt, are mean values ± s.e.m. (n=3). **B)** Expression levels of nNa_V_1.5, determined by Western Blot (WB) in breast cancer cell lines (MCF-7, SKBR3, MDA-MB-231, MDA-MB-468), EBNA-nNa_V_1.5. EBNA-nNa_V_1.5 as positive control for nNa_V_1.5. Cells were seeded onto FN for 2 hours. Representative WB (left panel) and densitometric analysis (right panel) are reported. Data are mean values ± s.e.m obtained in three independent experiments. a.u.= arbitrary units. **C)** Membrane expression levels of nNa_V_1.5, determined by Immunofluorescence (IF) in breast cancer cell lines (MCF-7, SKBR3, MDA-MB-231, MDA-MB-468). Cells were seeded onto FN for 2 hours. Cells were stained (in red) with mAb-nNa_V_1.5 primary Ab conjugated with AlexaFluor-647. Representative images (scale bar 100 µm) and quantification are reported. The quantification was performed considering only the membrane signal, highlighted by the white masks shown in the pictures. For each condition the fluorescence relative to 20 cells was analyzed**. For total IF signal, we measured the fluorescence intensity for each cell, dividing it by the cell area, to obtain a final mean fluorescence (± standard error mean).** Full images are reported in **Supplementary Figure S3B**. Pictures were taken on a confocal microscope (Nikon TE2000, Nikon; Minato, Tokyo, Japan). ImageJ software was used to analyze the images. **D)** Co-immunoprecipitation (co-IP) between hERG1, nNa_V_1.5 and β1 integrin in BCa cells (MCF-7, SKBR3, MDA-MB-231, MDA-MB-468) and HEK-293 (negative control) seeded onto FN for 2 hours. Immunoprecipitation was performed with monoclonal antibody recognizing extracellular epitopes of the β1 integrin subunit (TS2/16), and a monoclonal antibody recognizing hERG1 (mAb-hERG1), left panels. **Left panel:** representative experiment; **right panel:** densitometric analysis (n=3). Data are reported as a. u. (arbitrary unit). *P < 0.05 and **P < 0.01. IP = immunoprecipitation.

We therefore asked whether nNav1.5 could also be an additional component of the hERG1/β1 integrin macromolecular complex. Thus, co-IP (**Figure 3D**) and co-localization (**Figure 4A**) experiments were performed on all the BCa cells. For co-IP we used either the mAb-TS2/16 (IP: β1 integrin) or the mAb-hERG1 (IP: hERG1). The mAb-nNa_V_1.5 was then used in the WBs to detect the nNa_V_1.5 protein and the co-IPs were quantified by densitometry as detailed in Materials and Methods (**Figure 3D**). Both hERG1 and nNa_V_1.5 co-immunoprecipitated with β1 integrin, in different proportions in the cell lines (see the bar graph showing the densitometric data below **Figure 3D**). The two TNBCa cells, showed the highest expression levels of the three proteins. IF experiments, employing the two monoclonal antibodies (one directed against hERG1 and the other against nNa_V_1.5) and quantifying the co-localization as MOC, confirmed that hERG1 and nNa_V_1.5 co-localized in BCa cells seeded for 2 hours on FN (**Figure 4A**). Silencing either hERG1 or the β1 integrin not only led to a decrease of these proteins (Figure 4B and 4C) but also affected the expression of other proteins. Silencing hERG1 decreased the amount of nNa_V_1.5 (Figure 4B, lanes indicated as Input nNa_V_1.5) and its withdrawal from the co-IP (Figure 4B, lanes indicated as WB nNa_V_1.5). Silencing β1 integrin resulted in the complete withdrawal of both hERG1 and nNa_V_1.5 from the co-IP (Figure 4C, lanes indicated as WB hERG1 and WB nNa_V_1.5, respectively) but did not affect the expression of the two proteins (Figure 4C, lanes indicated as Input hERG1 and Input nNa_V_1.5, respectively). MDA-MB-468 cells showed the highest overlap (MOC = 0.76 ± 0.04) meaning the highest co-localization of the two channels. Finally, IF experiments also showed that nNa_V_1.5 co-localized with paxillin, meaning the presence of the channel in FA, as the hERG1/β1 integrin complex (**Figure 4D**).

**Figure 4.**
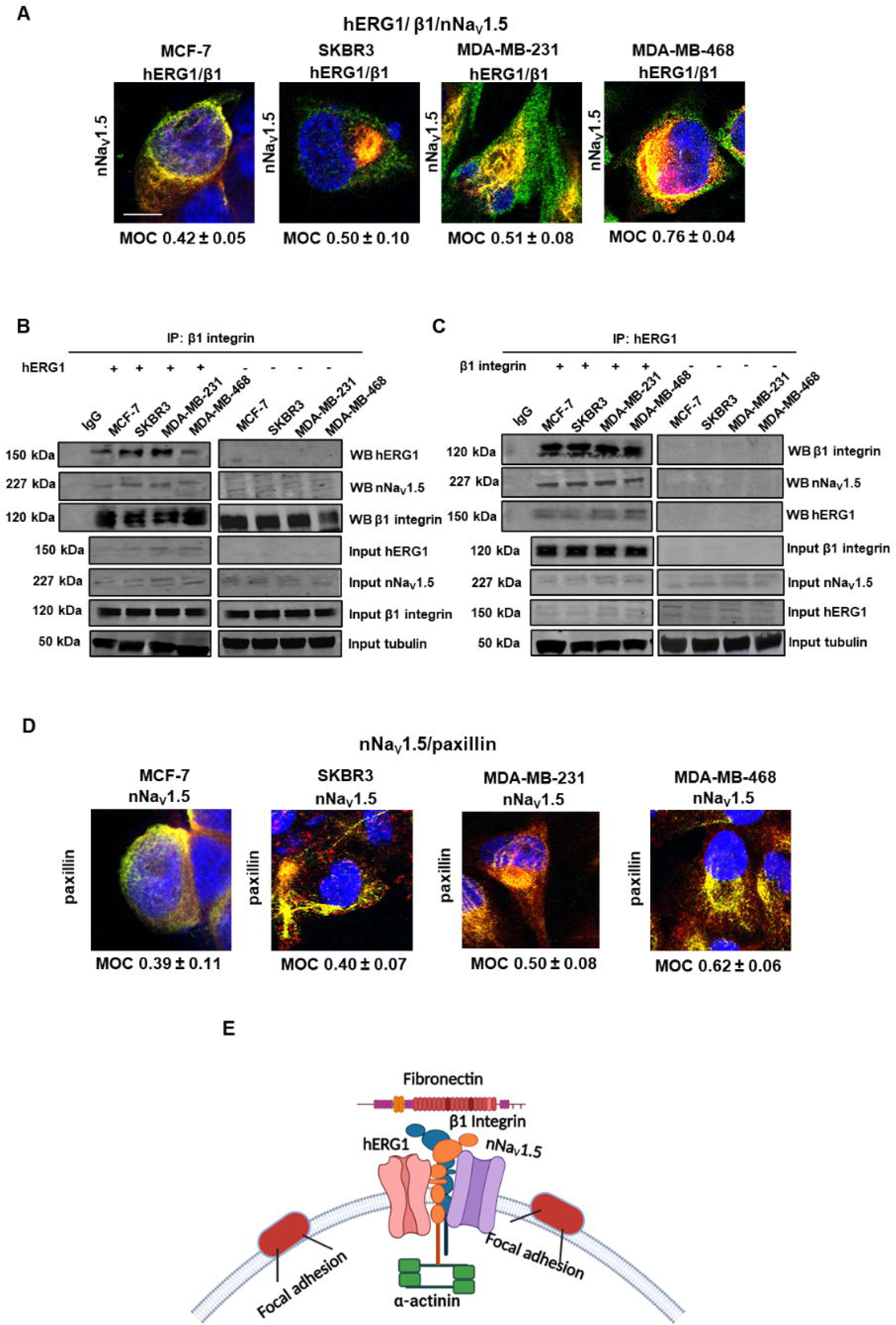
hERG1, β1, nNa_V_1.5 form a tri-molecular complex which is localized in focal adhesions. **A)** Representative IF images (scale bar: 100 μm) of the co-localization of hERG1, β1 integrin hERG1 (in green probed with scDb-hERG1-β1 conjugated with alexa-488) and nNa_V_1.5 (in red probed with mAb-nNa_V_1.5 conjugated with AlexaFluor-647) in breast cancer cell lines (MCF-7, SKBR3, MDA-MB-231, MDA-MB-468) seeded onto FN for 2 hours. Mander’s Overlapping Coefficient (MOC) reporting the proteins correlation is showed below each image. For each condition the fluorescence relative to 20 cells was analyzed. For total IF signal, we measured the fluorescence intensity for each cell, dividing it by the cell area, to obtain a final mean fluorescence (± standard error mean). Full images are reported in **Supplementary Figure S4A**. Pictures were taken on a confocal microscope (Nikon TE2000, Nikon; Minato, Tokyo, Japan). ImageJ software was used to analyse the images. **B)** Co-IP between hERG1, nNa_V_1.5, and β1 integrin in BCa transiently hERG1 and **C)** β1 integrin silenced cells (MCF-7, SKBR3, MDA-MB-231, MDA-MB-468) seeded onto FN for 2 hours. Immunoprecipitation was performed with monoclonal antibody recognizing hERG1 (mAb-hERG1) and monoclonal antibody TS2/16. Data are mean values ± s.e.m obtained in three independent experiments. a.u.= arbitrary units. **D)** Representative IF images (scale bar: 100 μm) of the co-localization of nNa_V_1.5 (in green probed with mAb-nNa_V_1.5 conjugated with AlexaFluor-488) and paxillin (in red probed with mAb-paxillin and secondary Ab conjugated with AlexaFluor-546) (bottom panel) in breast cancer cell lines (MCF-7, SKBR3, MDA-MB-231, MDA-MB-468) seeded onto FN for 2 hours. MOC reporting nNa_V_1.5/paxillin correlations is shown below each image. For each condition the fluorescence relative to 20 cells was analyzed. For total IF signal, we measured the fluorescence intensity for each cell, dividing it by the cell area, to obtain a final mean fluorescence (± standard error mean). Full images are reported in **Supplementary Figure S4B**. Pictures were taken on a confocal microscope (Nikon TE2000, Nikon; Minato, Tokyo, Japan). ImageJ software was used to analyse the images. **E)** Model representation of the macromolecular complex comprising hERG1, β1 integrin, α-actinin and nNa_V_1.5. Created with BioRender.com. IP = immunoprecipitation. MOC = Manders’ Overlapping Coefficient.

In conclusion, a tri-molecular complex hERG1/β1/nNa_V_1.5 occurs in FAs of BCa cells, especially TNBCa cells, once seeded on FN for 2 hours (**Figure 4E**).

### hERG1 and nNa_V_1.5 are mutually “controlled” in BCa cells

Since hERG1 and Na_V_1.5 form a microtranslatome in the heart^33^, we tested whether the two channels were linked at the translational level also in BCa cells. To this purpose, we analyzed the effects of either silencing the two channels separately or overexpressing one of the two channels, in particular overexpressing hERG1^15^. As already observed from data shown in **Figure 4B**, silencing hERG1 induced a decrease in nNa_V_1.5 expression by 40-45%, in MCF-7 and SKBR3, and by 55% ca in TNBCa cell lines, in addition to the disappearance of the hERG1 protein (**Figure 5A**). Silencing nNa_V_1.5 resulted in the disappearance of the nNa_V_1.5 protein, accompanied by a decrease of hERG1 expression by 45% in both MCF-7 and SKBR3, and by 55% ca. in the TNBCa cell lines (Figure 5B). IF experiments silencing nNa_V_1.5 show that hERG1 (probed with mAb-hERG1) is undetectable on the plasma membrane in MDA-MB-231, as reported in **Supplementary Figure S5A**. These results are consistent with the fact that it’s impossible to measure any hERG1 current.

**Figure 5.**
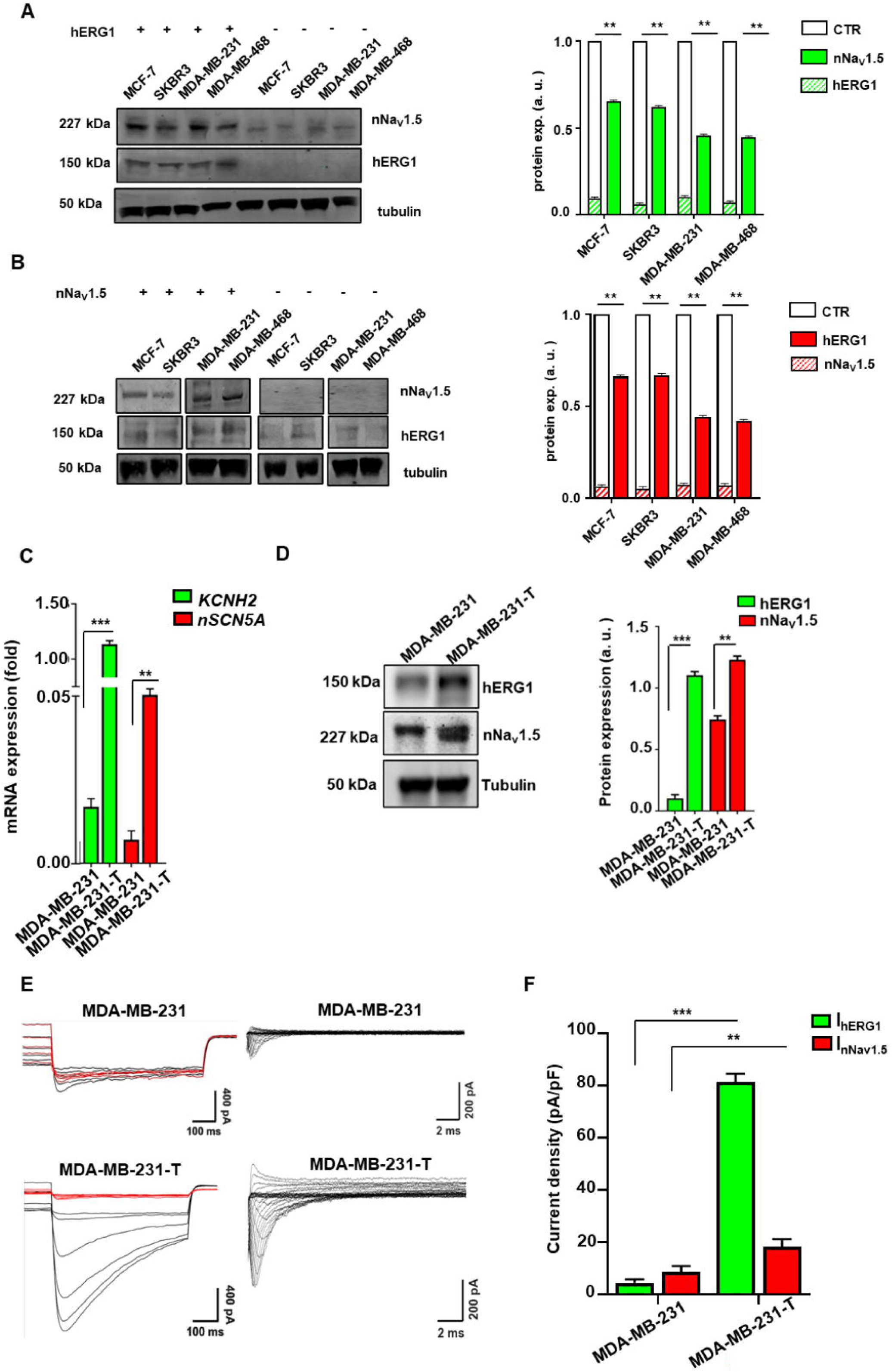
The expression of hERG1 and nNa_V_1.5 is mutually regulated in BCa cells. WB revealing hERG1, nNa_V_1.5 and β1 integrin in BCa transiently hERG1 (**A)** and nNa_V_1.5 (**B**) silenced cells (MCF-7, SKBR3, MDA-MB-231, MDA-MB-468) seeded onto FN for 2 hours. Left panel: representative experiment; right panel: densitometric analysis. Data are mean values ± s.e.m obtained in three independent experiments. a.u.= arbitrary units. **C)** Expression levels of the two transcripts, *KCNH2* and *nSCN5A*, determined by Real Time PCR in MDA-MB-231 and MDA-MB-231-T cell lines. Data, reported as 2-DCt, are mean values ± s.e.m. (n=3). **D)** Expression levels of the two proteins, hERG1 and nNa_V_1.5, determined by WB in MDA-MB-231 and MDA-MB-231-T cell lines. Representative WB (left panel) and densitometric analysis (right panel) are reported. Data are mean values ± s.e.m obtained in three independent experiments. **E)** On the left, patch clamp hERG1 inward current trace and activation curve of a representative MDA-MB-231 and MDA-MB-231-T cell. Patch clamp hERG1 inward current trace recorded in MDA-MB-231 cells shown below it. On the right Na^+^ traces recorded in MDA-MB-231 and MDA-MB-231-T cells. Quantitative analysis is shown on the right. Data are mean values ± s.e.m obtained in three independent experiments. For all experiments of lateral motility and invasiveness, a background level of invasion was calculated, seeding cells without chemotactic gradient, and a basal level of invasion was then subtracted from the experiment measurements. **P < 0.01 and ***P < 0.001. IP = immunoprecipitation. WB = Western Blot.

The over-expression of *KCNH2* in MDA-MB-231 cells (giving MDA-MB-231-T cells) produced a significant over-expression of both *KCNH2* and *nSCN5A* transcripts (**Figure 5C**). These results were also observed at the protein level (WB data are in **Figure 5D**, IF data are in **Supplementary Figure S5B**) as well as in the recorded specific currents (**Figure 5E**) ^17,34,35^. Indeed I_hERG1_ was 98.8 ± 26.4 pA (N = 5 cells) in MDA-MB-231 WT and 2673.0 ± 143.0 pA (N = 5 cells) in MDA-MB-231-T. I_nNaV1.5_ was 172.8 ± 20.2 pA (N = 5 cells) in MDA-MB-231 WT and 478.2 ± 69.8 pA (N = 8 cells) in MDA-MB-231-T. Cells from both cell lines were then recorded in control condition and after E4031 (1µM) 1 min application (**Supplementary Figure S5C**). Their features appeared to be compatible with our previous models (HEK-hERG1 and SH-SY5Y; V_1/2_ (mV): −54.5 ± 2.3), as shown in **Supplementary Figure S5D**^36^. The respective average current densities are reported in **Figure 5F**. Overall, the hERG1 and nNa_V_1.5 channels are mutually controlled in BCa cells, at both the transcription and translation level, similarly to what occurs in the heart^33^. This result was further stressed analyzing the effects of the two specific blockers of either hERG1 (i.e. E4031) or Na_V_1.5 (i.e. TTX). E4031 was used at 40 μM concentration as in Masi et al^37^ and in Duranti and Iorio et al^10^; TTX was used at 20 μM concentration according to Brackenbury et al^19^. The treatment with E4031 decreased the expression of both hERG1 and nNa_V_1.5 by 45% ca. in MCF-7 and SKBR3 and by 55%-60% in TNBCa cell lines (**Figure 6A**). The treatment with TTX decreased the expression of both nNa_V_1.5 and hERG1 by 40% in both MCF-7 and SKBR3 and by 55%-60% in the TNBCa cell lines (**Figure 6B**). Overall, the two channel blockers decreased the expression of the two channel proteins with a stronger effect in the TNBCa cell lines (see the bar graphs with the densitometric data on **Figure 6C**). Finally, both E4031 and TTX decreased the level of the hERG1/β1/nNa_V_1.5 complex to the same extent, although a clear trend it’s shown where E4031 decreases the complex more than TTX. In addition to that the effect is clearly higher in the two TNBCa cell lines (**Figure 6D** and the bar graphs with the densitometric data in **Figure 6E**).

**Figure 6.**
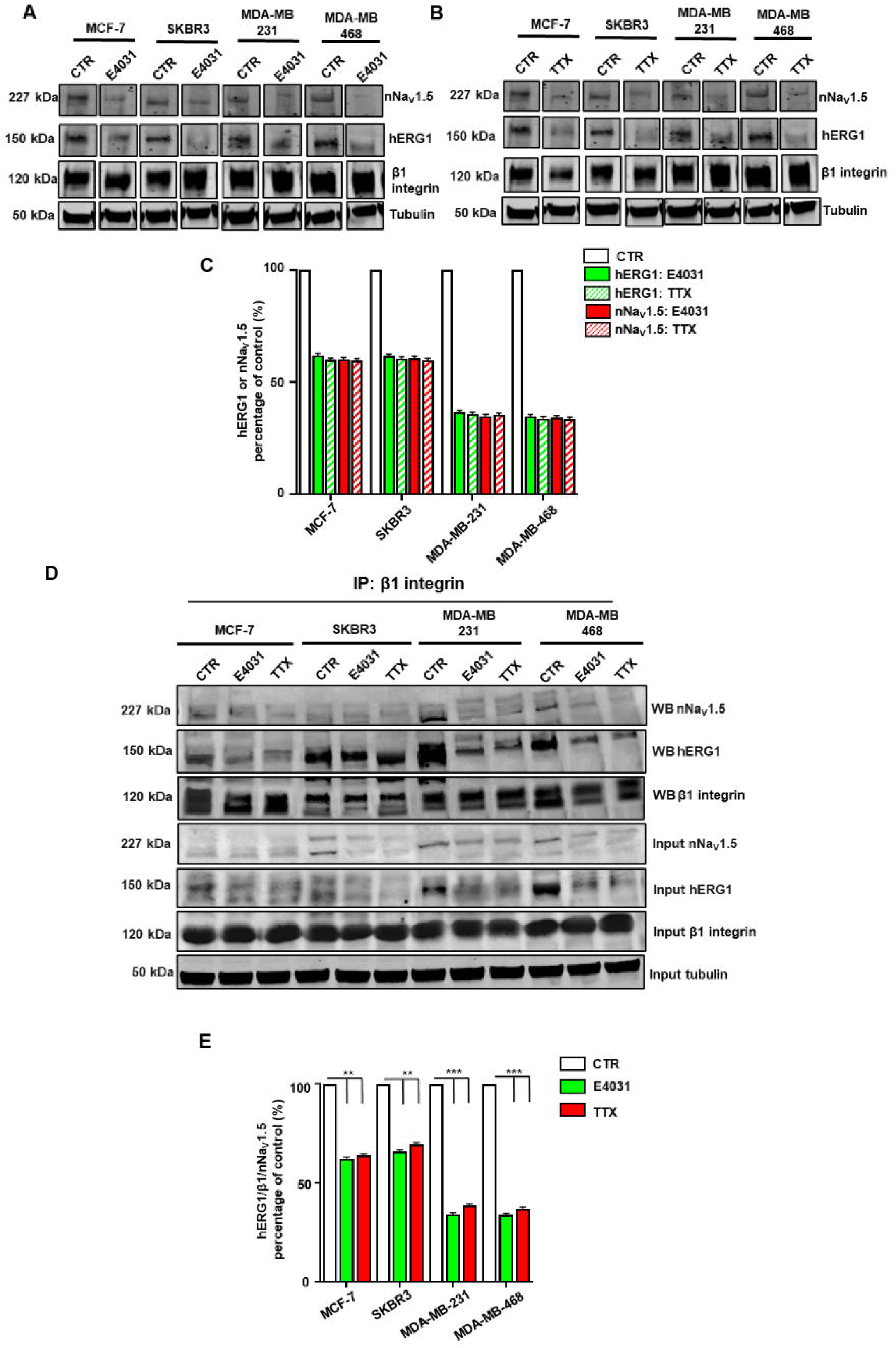
hERG1 and nNa_V_1.5 form a macromolecular complex with β1 integrins which is inhibited by either hERG1 or Na_V_ blockers. **A)** WB of all TNBCa cell lines treated with E4031 (40 μM) and **B)** with TTX (20μM) with the respective densitometric analysis **C)** in panel below. All the quantification were normalized against control (not shown). **D)** co-IP between hERG1, nNa_V_1.5 and β1 integrin in BCa cells (MCF-7, SKBR3, MDA-MB-231, MDA-MB-468) seeded onto FN for 2 hours and treated with E4031 (40μM) and TTX (20μM). Immunoprecipitation was performed with monoclonal antibody TS2/16. **E)** Densitometric analysis. Data are mean values ± s.e.m obtained in three independent experiments. a.u.= arbitrary units. **P < 0.01 and ***P < 0.001. IP = immunoprecipitation. WB = Western Blot.

### The hERG1/**β**1/nNav1.5 complex controls f-actin organization

Based on the role of the hERG1/β1 integrin complex in PDAC and CRC cells ^10^, we then studied the involvement of the hERG1/β1 integrin complex in f-actin organization of BCa cells. To this purpose, Bca cells were seeded on FN for two hours, fixed and processed for IF to determine (i) the expression and recruitment of the Arp2/3 complex (i.e. the first step of f-actin enucleation ^38,39^; (ii) the organization of f-actin in stress fibers and filopodia, using rhodamine conjugated phalloidin as in^10^; (iii) the emission and quantification of invadopodia, staining the cells with an anti-cortactin antibody^40^. To determine the role of the complex, we treated BCa cells with E4031, TTX or the scDb-hERG1-β1, since all these treatments reduced complex formation (see Figure 1E, 1F and 5D), although the effect of scDb-hERG1-β1 was stronger.

The fluorescence intensity of Arp2/3 significantly increased at the peripheric areas of cell surface (see the white masks in Figure 2C) when MDA-MB-231 and MDA-MB-468 were treated with E4031, and even more with TTX or scDb-hERG1-β1 compared to controls (**Figure 7A**). This indicates that the Arp2/3 complex moves from the cytoplasm to peripheric areas, possibly those where amoeboid protrusions (filopodia) are formed (see below). No difference in the localization of Arp2/3 was seen in MCF-7 and SKBR3 when treated with either E4031, TTX or scDb-hERG1-β1 (**Supplementary Figure S7D**). Both E4031 and TTX reduced the length of stress fibers by roughly 50% and 55%, respectively. The treatment with scDb-hERG1-β1 had an even stronger effect (**Figure 7B** and the bar graph on the bottom). The length and density of filopodia were also increased by all the treatments (**Figure 7B** and the bar graph on the bottom). TTX and scDb-hERG1-β1 had the strongest effect (30-40%). A similar increase was observed by all the treatments on the density of cortical actin (**Figure 7B** and the bar graph on the bottom). Even in this case TTX and scDb-hERG1-β1 had the strongest effect (40-45%). The staining with the anti-cortactin antibody showed a clear, punctate pattern in the two TNBCa cell lines (**Figure 7C**). In contrast, cortactin staining was weak and diffuse and only few, faint spots were detected in both MCF-7 and SKBR3 cells (**Supplementary Figure S7E**). The Invadopodia Index (InvI) value was reduced by E4031 (roughly 55%) and even more by TTX (65%) and scDb-hERG1-β1 (roughly 60%) in MDA-MB-231 and MDA-MB-468 cells (**Figure 7C** and the bar graph on the right). No significant effect was seen in MCF-7 and SKBR3 (**Supplementary Figure S7F**).

**Figure 7.**
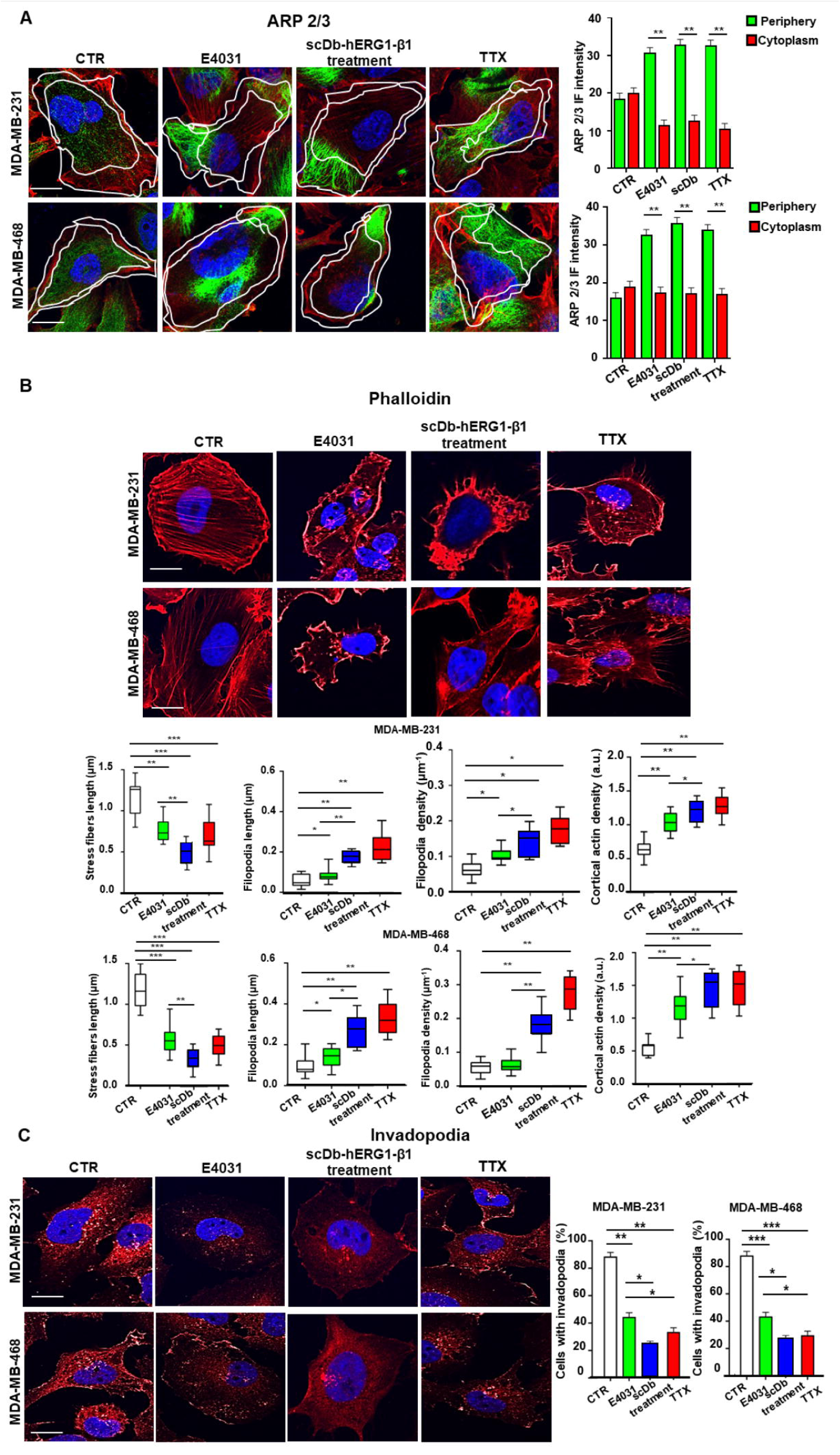
hERG1 and nNa_V_1.5 modulates the organization of f-actin. **A)** Staining with anti-Arp2/3 (green) antibody with secondary Ab AlexaFluor-488 and CellMask™ Deep Red Actin Tracking Stain (red) of MDA-MB-231 and MDA-MB-468 cells seeded on FN for 2 hours and treated with E4031: 40μM, scDb-hERG1/β1: 20µg/ml and TTX: 20μM. Corresponding Arp2/3 graphs on the right panel. **B)** Phalloidin staining (Ab conjugated with Rhodamine) of MDA-MB-231 and MDA-MB-468 cells seeded on FN for 2 hours and treated with E4031: 40μM, scDb-hERG1/β1: 20µg/ml and TTX: 20μM. Corresponding graphs of actin stress fibers length, filopodia length and cortical actin density. **C)** Staining with anti-cortactin antibody of MDA-MB-231 and MDA-MB-468 cells seeded on FN for 2 hours and treated with E4031: 40μM, scDb-hERG1/β1: 20µg/ml and TTX: 20μM. Corresponding graphs of the percentage of cells that show invadopodia. At least 20 cells per condition were analyzed and all p-values were determined by a Mann–Whitney test for non-parametric values, or for data deviating from normality by a Kolmogorov–Smirnov test. Scale bars: 10 μm. Full images for all panels are reported in **Supplementary Figure S7A, B and C**. Pictures were taken on a confocal microscope (Nikon TE2000, Nikon; Minato, Tokyo, Japan). ImageJ software was used to analyse the images. In the graphs, boxes include central 50% of data points, the horizontal lines denote minimum value, median and maximum value. Data are mean values ± s.e.m. (*n*=3). *P < 0.05, **P < 0.01 and **P < 0.001. IF = immunofluorescence. FN = Fibronectin.

Overall, these data indicate that the hERG1/β1/nNa_V_ 1.5 integrin complex regulates the organization of f-actin in TNBCa cells, keeping the stress fibers spread and neatly grouped, and the filopodia retracted, to allow the organization of f-actin in invadopodia.

### The signaling pathways downstream to the hERG1/**β**1/nNa_V_1.5 complex

We then studied the intracellular signaling pathways which is triggered by the hERG1/β1/nNav1.5 complex and regulates f-actin organization in the different structures of the cytoskeleton. We first focused on (i) intracellular Ca^2+^ ^41^ and (ii) AKT^10,15^. These experiments were performed on MDA-MB-468 (which had the highest levels of the complex and on MCF-7 cells which showed the lowest (see Figure 1D and 3D). The basal value of the intracellular calcium concentration ([Ca^2+^]_i_) was significant higher in MDA-MB-468 cells compared to MCF-7 (Figure 7A), but no effect of plating on FN or treating the cells with either channel blocker in either cell line was observed (**Figure 8A**). No effects in either cell lines treating with either channel blockers or the scDb-hERG1-β1 was also observed in the levels of AKT phosphorylation (pAKT), as determined by both WB (**Figure 8B**) and IF (**Supplementary Figure S8A&B**).

**Figure 8.**
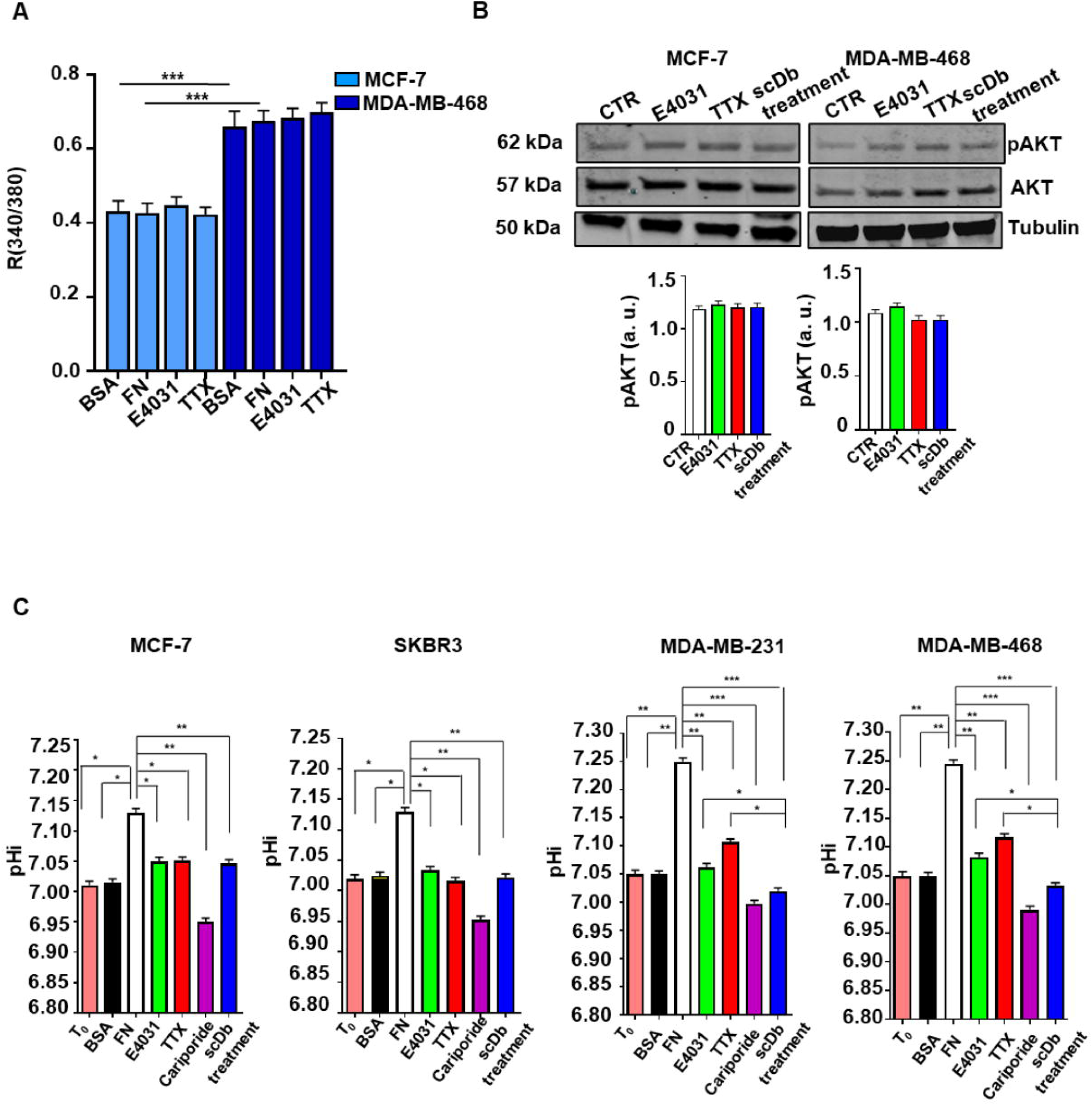
The signaling pathways downstream to the hERG1/β1 integrin/nNa_V_1.5 complex. **A)** The intracellular Ca^2+^ has been also measured under different conditions. MCF-7 and MDA-MB-468 have been seeded on different type of coating method: FN and BSA for 2h and then intracellular Ca^2+^ was measured as reported in Materials and Methods. Image acquisition (3s frequency) was performed using Metafluor Imaging System (Molecular Devices; Sunnyvale, CA, USA). **B)** pAKT levels were determined by Western Blot (WB) in MCF-7 and MDA-MB-468 while cells were treated with specific blockers E4301 (40µM), TTX (20µM) and scDb-hERG1-β1 (20µg/ml) in different coating conditions (BSA and FN for 2h). Representative WB (top panel) and densitometric analysis (bottom panel) are reported. Data are mean values ± s.e.m obtained in three independent experiments. a.u.= arbitrary units. **C)** pH_i_ values of MCF-7, SKBR3, MDA-MB-231 and MDA-MB-468 cell lines seeded for 2h onto FN and treated with E4031 (40μM), TTX (20μM), cariporide (5μM), scDb-hERG1-β1 (20µg/ml) and Ouabain (300nM). All pH values were calculated using 490/440nm fluorescence ratio and applying standard curve and linear equations. Data are mean values ± s.e.m obtained in three independent experiments. *P < 0.05, **P < 0.01 and ***P < 0.001. R = ratio.

Given the relevant role of pH (both extracellular, pHe, and intracellular, pHi) on f-actin organization and hence cancer cell migration and progression^42–44^ and the role of pHi in the hERG1-dependent regulation of cell migration in colorectal cancer cells^11^, we studied pHi in our model. Compared with the initial value (T0), pH_i_ increased significantly in all the cells when plated on FN for 2h (**Figure 8C**, pink vs white bars): 7.013 ± 0.002 to 7.135 ± 0.001 (in MCF-7 cells), 7.042 ± 0.001 to 7.171 ± 0.002 (in SKBR3 cells), 7.052 ± 0.003 to 7.253 ± 0.002 (in MDA-MB-231 cells) and 7.081 ± 0.001 to 7.283 ± 0.003 (in MDA-MB-468 cells). Thus, the FN-induced intracellular alkalinization was higher (by ca. 15%) in the TNBC cells. No change was seen in cells seeded on BSA control (**Figure 8C**, black bars). E4031 and TTX equally reduced the FN-induced alkalinization of pH_i_, especially in the TNBC cells, while scDb-hERG1-β1 had a significantly stronger effect. We then tested whether the Na^+^/H^+^ antiporter NHE1 was involved in the above pHi alkalinization. Indeed, the specific NHE1 inhibitor cariporide completely blocked the effect of cell adhesion to FN on pHi, decreasing pH_i_ to levels lower than those measured at T0 or in cells seeded on BSA (**Figure 8C**). Overall, NHE1 is the main responsible of pHi regulation in BCa cells, and in particular in the FN-induced alkalinization observed in TNBCa.

We then investigated whether NHE1 was present in the hERG1/β1/nNa_V_1.5 macromolecular complex. Indeed, the antiporter co-immunoprecipitated with β1 integrin, hERG1 and nNa_V_1.5 (**Figure 9A and B**). Interestingly the antiporter was absent when hERG1 was transiently silenced, indicating its binding to the K^+^ channel protein (**Figure 9A)**. On the other hand, silencing β1 did not result in the absence of NHE1 showing that its binding is not related to the β1 integrin (**Figure 9B**). Hence, the complex can be addressed as “NHE1/hERG1/β1/nNaV1.5 complex. The presence of NHE1 antiporter in the complex turned out to be relevant in the FN-induced f-actin organization, since the treatment with cariporide decreased stress fiber length and increased filopodia length (**Figure 9C**), similarly to what was observed with the E4031, scDb-hERG1-β1 and TTX treatments. Again, these effects were stronger in TNBCa cells MDA-MB-231 and MDA-MB-468 (**Figure 9D**; **Supplementary Figure S9A**).

**Figure 9.**
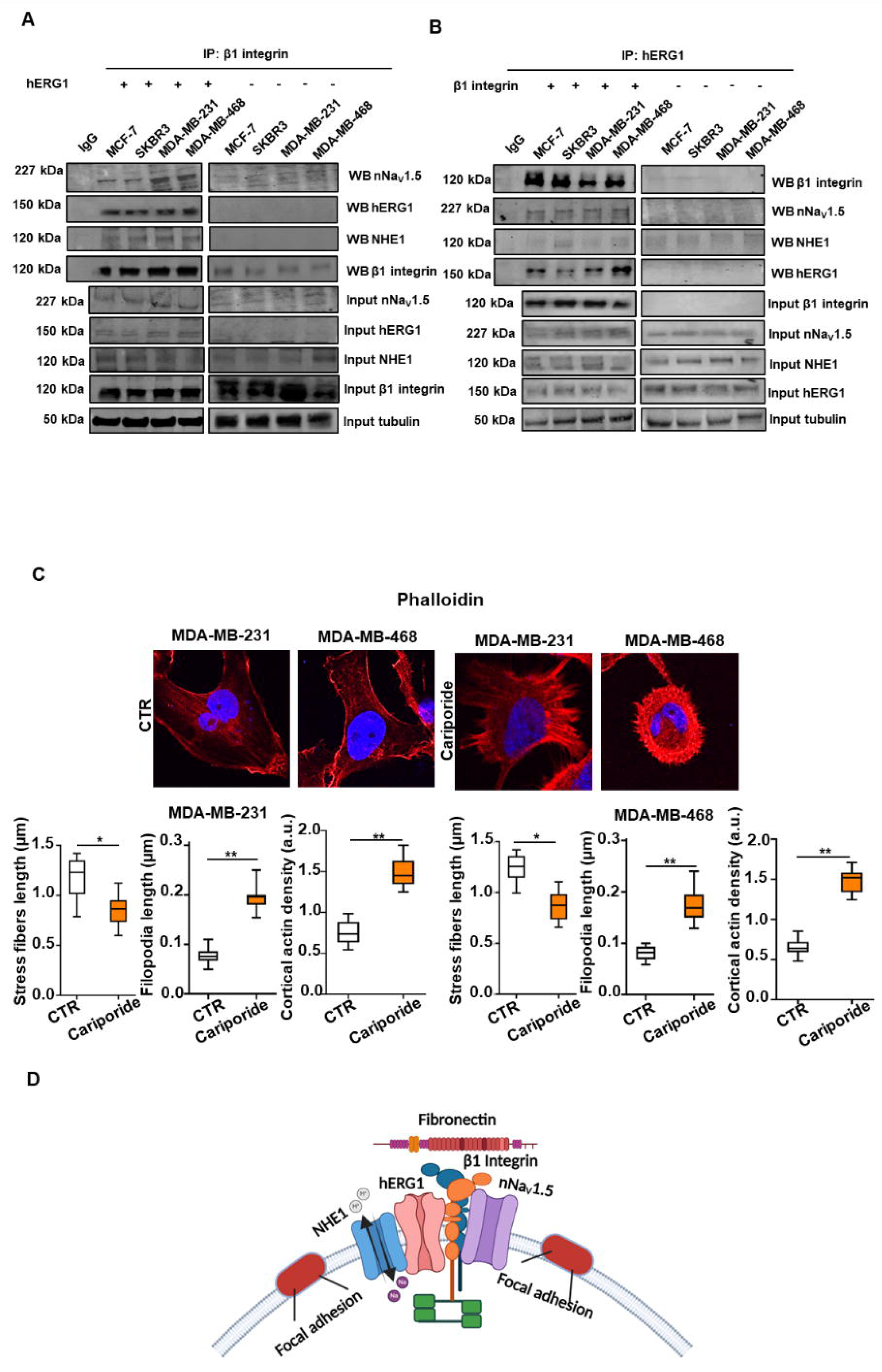
The hERG1/β1/nNa_V_1.5 complex comprises the Na^+^/H^+^ exchanger NHE1. Effects of the NHE1 inhibitor cariporide on f-actin organization in TNBCa cells. **A)** co-IP between hERG1, nNa_V_1.5, NHE1 and β1 integrin in BCa transiently hERG1 and **(B)** β1 silenced cells (MCF-7, SKBR3, MDA-MB-231, MDA-MB-468) seeded onto FN for 2 hours. Immunoprecipitation was performed with monoclonal antibody TS2/16 (A) and monoclonal antibody recognizing hERG1 (mAb-hERG1) (B). In the graphs, boxes include central 50% of data points, the horizontal lines denote minimum value, median and maximum value. Data are mean values ± s.e.m. (*n*=3). **C)** Phalloidin (conjugated with rhodamine) staining of MDA-MB-231 and MDA-MB-468 cells seeded on FN for 2 hours and treated with Cariporide (5μM). Representative images are on the top part of the panel, the corresponding graph of actin stress fibres length, filopodia length and density are below. At least 20 cells per condition were analyzed and all p-values were determined by a Mann–Whitney test, or for data deviating from normality by a Kolmogorov–Smirnov test. Scale bars: 10 μm. Full images are reported in **Supplementary Figure S9A**. Pictures were taken on a confocal microscope (Nikon TE2000, Nikon; Minato, Tokyo, Japan). ImageJ software was used to analyse the images. **D)** Model representation of the macromolecular complex comprising hERG1, β1 integrin, α-actinin, nNa_V_1.5 and NHE1. Created with BioRender.com. IP = immunoprecipitation. *P < 0.05, **P < 0.01.

Overall, these data show that a NHE1/hERG1/β1/nNa_V_1.5 regulates pHi and affects f-actin organization in TNBCa cells (**Figure 9D**).

### The hERG1/β1 integrin/nNa_V_1.5 complex promotes TNBCa cell migration and invasiveness

We then analyzed the contribution of the NHE1/hERG1/β1/nNav1.5/NHE1 complex to cell proliferation, lateral motility and Matrigel invasion of the four cell lines We tested the effects of (i) E4031, (ii) TTX and (iii) scDb-hERG1-β1 which similarly impaired the formation of the complex. Neither treatment had any effect on proliferation (**Figure 10A**). In contrast, both E4031 and TTX significantly inhibited the lateral motility of MDA-MB-468 and MDA-MB-231 cells, reducing MoI by ca. 50% after 24h (**Figure 10B**). The effect of scDb-hERG1-β1 was even stronger compared to the two blockers reducing MoI by roughly 55%. Cariporide reduced cell motility (a reduction of MoI of 75% was observed), and the strongest effect (reduction of MoI of 85%) was obtained by the combination of Cariporide and scDb-hERG1-β1. The motilities of the MCF-7 and SKBR3 cells were only slightly or not affected by E4031, TTX, scDb-hERG1-β1, cariporide or the combination of cariporide and scDb-hERG1-β1 (**Figure 10B**). Similar observations were made in experiments lasting 48h (**Supplementary Figure S10A**).

**Figure 10.**
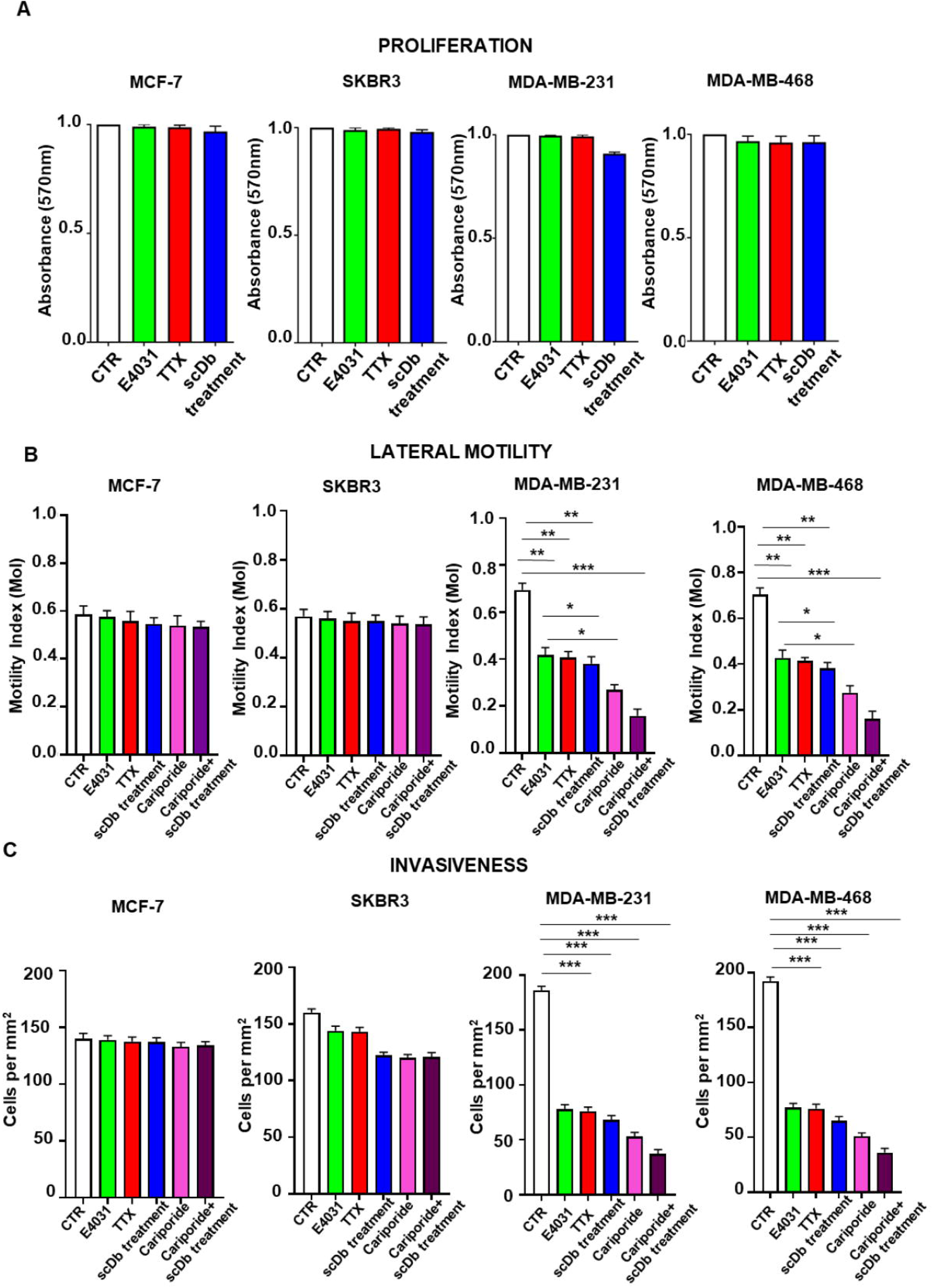
hERG1 and nNa_V_1.5 control cell motility and invasiveness in BCa cell lines. **A)** Cell proliferation determined by MTT test in MCF-7, SKBR3, MDA-MB-231 and MDA-MB-468 and cell lines. Cells were treated with E4031 (40μM), tetrodotoxin (TTX) (20μM) and scDb-hERG1-β1 (20µg/ml) for 24 hours and absorbance at 570n was measured. **B)** Motility Index (MoI) determined by lateral motility experiments in MCF-7, SKBR3, MDA-MB-231 and MDA-MB-468 cell lines. Cells were treated with E4031 (40μM), tetrodotoxin (TTX) (20μM), scDb-hERG1-β1 (20µg/ml), cariporide (5µM) and a combination of the latters. Data are mean values ± s.e.m obtained in three independent experiments. **C)** Invasiveness obtained with matrigel invasion assay in MCF-7, SKBR3, MDA-MB-231 and MDA-MB-468 cell lines. Cells were treated with E4031 (40μM), tetrodotoxin (TTX) (20μM), scDb-hERG1-β1 (20µg/ml), cariporide (5µM) and a combination of the latters for 24 hours and cells per mm^2^ were counted. Data are mean values ± s.e.m obtained in three independent experiments. The number of cells counted in each field of view was converted to “cells per mm^2^” by multiplying the number of counted cells by a factor of 4.07. For all experiments of lateral motility and invasiveness, a background level of invasion was calculated, seeding cells without chemotactic gradient, and a basal level of invasion was then subtracted from the experiment measurements. *P < 0.05, **P < 0.01 and **P < 0.001.

**Figure 11.**
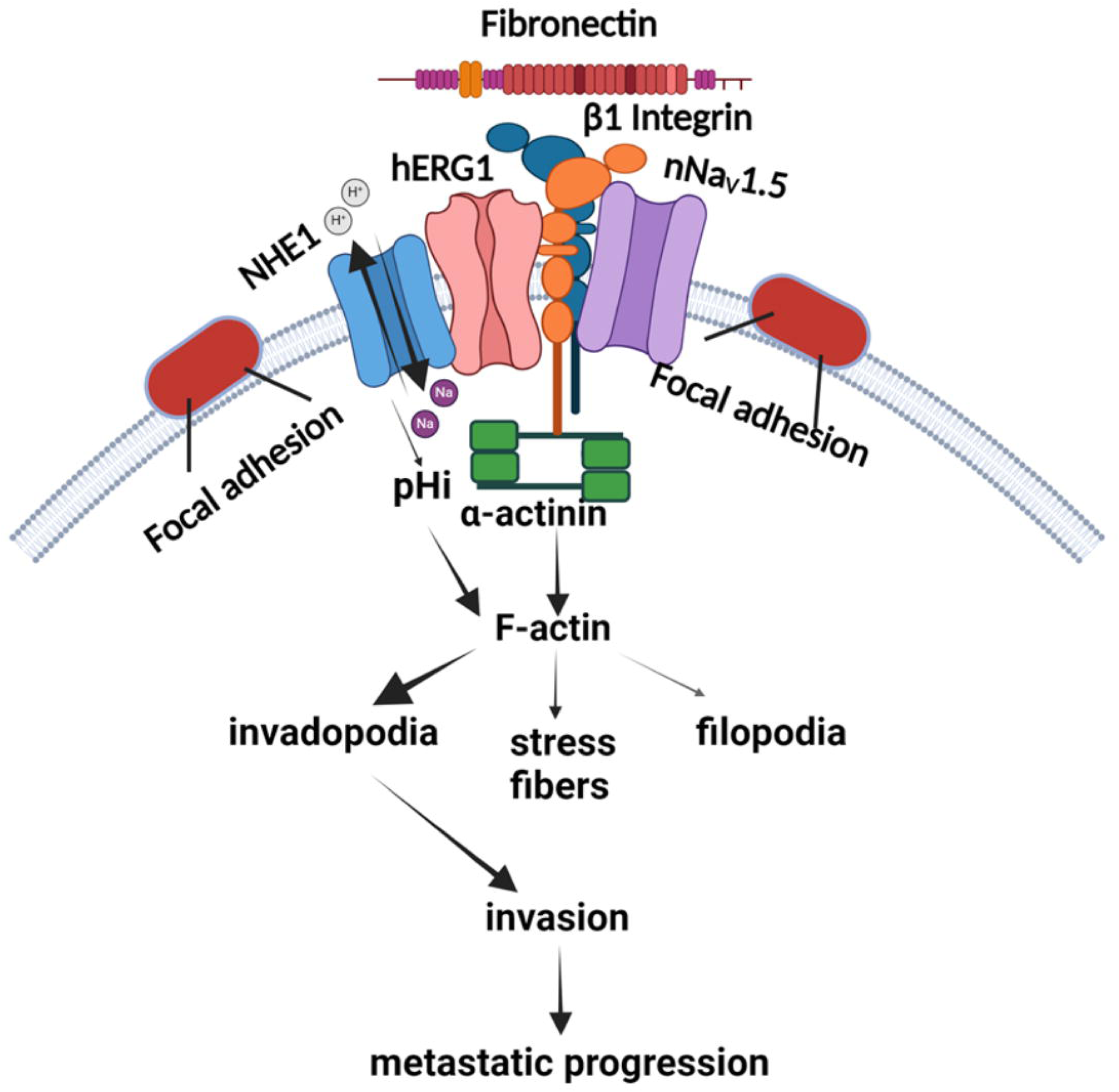
The integrin-centered macromolecular signaling complex in BCa cells. In cells adhering to FN a macromolecular complex is formed which comprises hERG1, the β1 integrin subunit and the neonatal form of Na_V_1.5. The β1 integrin associates with α-actinin which directly modulates f-actin organization. hERG1 also associates with the NHE-1 exchanger whose activation increases the intracellular pH and impacts on f-actin organization. The f-actin remodeling leads to the formation of invadopodia. Created with BioRender.com.

As regards Matrigel invasion, E4031, TTX, scDb-hERG1-β1 and cariporide were significantly inhibitory on MDA-MB-231 and MDA-MB-468 cells (ca. 40% and 40%; 40% and 45%; 45% and 45%; 50% and 50%; 60% and 60% respectively) after 16h (**Figure 10C**). Also in this case, the combination of cariporide and scDb-hERG1-β1 showed the strongest inhibitory effect on Matrigel invasion. Only a slight or no effect were observed on SKBR3 and MCF-7 cells with all the above treatments.

## DISCUSSION

Cancer metastases are the main obstacle to successfully treating solid cancer, especially hard-to-treat-cancers such as Triple Negative Breast Cancer (TNBCa). The pro-metastatic behavior of malignant cells is regulated by signals produced by the Tumor Microenvironment (TME), often as a consequence of integrin-mediated cell adhesion to ECM proteins, which triggers pro-migratory, and hence pro-metastatic signaling pathways. In this context, integrin receptors coordinate signaling hubs constituted by multiprotein plasma membrane complexes often comprising ion channels and transporters. In the present paper, we uncovered a novel signaling pathway specifically triggered by the activation of β1 integrins in BCa cells seeded onto the ECM protein FN. This pathway, which occurs in BCa cells and in particular in TNBCa, involves a β1 integrin-centered plasma membrane macromolecular complex comprising the hERG1 K^+^ channel, the nNa_V_1.5 Na^+^ channel, the NHE1 Na^+^/H^+^ transporter and the cytoskeletal actin-binding protein α-actinin1. The complex activation by cell adhesion stimulates f-actin organization and thus TNBCa migration and invasiveness both directly (through α-actinin1 engagement) and indirectly (by NHE1-mediated cytoplasmic alkalinization). The expression of the two ion channels present in the complex is mutually regulated and their physiological role is essential in modulating the downstream signals and their functional consequences. The latter are in fact inhibited by using specific channel blockers, such as E4031 and TTX, and by RNA silencing procedures. In particular, hERG1 appears to exert a major structural role in the membrane complex, irrespective of hERG1 current activation. This is suggested by the observation that the downstream signaling is inhibited by either harnessing the hERG1/β1 integrin complex through a single chain bispecific antibody (scDb-hERG1-β1)^22^, or treating the cells with E4031. In fact, scDb-hERG1-β1 disrupts the hERG1/β1 integrin macromolecular complex without blocking the current^10^, whereas E4031 impairs the complex formation by blocking the channel in the open state, which has a lower affinity for β1 integrin^15^. Na_V_1.5 is also essential for downstream signaling, as is testified by our results with TTX. However, the balance of the contributions given by Na^+^ current activation and non-conductive Na_V_1.5 functions remains to be determined.

### The signaling pathway triggered by the hERG1/**β**1/nNa_V_1.5 macromolecular complex: pH_i_ regulates f-actin organization

The macromolecular complex we found in BCa cells is centered on the β1 subunit of integrin receptors, is activated by cell adhesion to the ECM protein FN and is localized in nascent focal adhesions. These features highlight the TME role in regulating BCa cell behavior, and particularly the aberrant migration that underlies cancer metastasis^45^. Interestingly, the macromolecular complex recruits NHE1 and α-actinin. These proteins are known to regulate cell migration^46^. The non-muscle α-actinin isoforms are actin-binding proteins that are necessary for proper cytoskeleton organization. In addition, α-actinins are associated with focal contacts and stress fibers and serve as scaffold proteins which connect the plasma membrane to the actin cytoskeleton and a variety of signaling pathways^47^. In BCa cells, α-actinin1 is bound to β1 integrins^30^ and its expression is elevated in TNBCa cell lines where it is involved in the rearrangement of actin cytoskeleton and the ensuing cell migration^48^. Furthermore, α-actinin is a direct target of the integrin-dependent Focal Adhesion Kinase (FAK) in platelets^49^ and is highly sensitive to pH values higher than 7^50^. When phosphorylated by FAK or in the presence of an alkaline pH, α-actinin loses its actin binding property^51^, which strongly enhances cell motility^47^. All these conditions are effective in our cellular model. In addition, NHE1 activity affects both pHi and pHe leading to alkalinization of pHi and acidification of pHe. This situation characterizes cancer cells^52^ and modulates their behavior, especially migration^53^. In particular, Na_V_1.5 has been proposed to promote MDA-MB-231 BCa cell migration and invasion through NHE1-dependent peri-membrane acidification, in turn stimulating acidic cathepsin^54,55^. Interestingly, Na_V_1.5 and NHE1 co-localize in caveolin-1-rich plasma membrane domains^31^, which further stresses the functional relevance of macromolecular plasma membrane complexes in the control of cellular pH. In our model, however, the effects of pH are mainly related to the organization of f-actin cytoskeleton. F-actin remodeling is known to determine different subcellular structures (filopodia, lamellipodia, and stress fibers), regulated by different signaling pathways^56^. In our system, the intracellular signals sustained by engagement of the NHE1/hERG1/β1/nNa_V_1.5 complex apparently converge to keep TNBC cells in a state readier to start the migration process.

The occurrence and functional relevance of ion channels within plasma membrane macromolecular complexes is emerging in both normal and pathological contexts^33,57–59^. An intriguing difference between ion channel complexes in normal vs cancer cells is the presence in the latter of interacting proteins that are relevant for cancer signaling^60,61^. Besides the hERG1/β1 integrin complex^15,62^, we notice that Orai1/TRPC1/SK3^63^, K_V_10.1/Calmodulin^64^, Kv10.1/Orai1/SPCA2^65,66^ and SWELL (LRRC8A)/integrin^67^ complexes have also been discovered in cancer cells. These macromolecular structures participate in different aspects of tumor behaviour that depend on cell adhesion to ECM proteins^68–70^.

### hERG1 and nNa_V_ 1.5 channels are co-translated and functionally cooperate in TNBCa

We notice that the two channel types present in the β1 integrin-centered complex in BCa cells, hERG1 and nNa_V_1.5, are also expressed in heart myocytes^71^. To warrant proper functional interplay in shaping the cardiac action potential, their expression is coordinated during protein translation through the hERG1/Na_V_1.5 “microtranslatomes”^33,72^. This observation agrees with the present data from BCa cells in which the two channel types were either silenced (or pharmacologically blocked) or transfected with hERG1. In particular, overexpressing hERG1 led to increased nNa_V_1.5 expression, as determined by quantifying mRNA, protein and current levels. These results suggest that the ratio between the amounts of these channel types needs to be precisely calibrated also in non-excitable cells such as BCa cells. We hypothesize that the balance of hERG1 and Na_V_1.5 can affect key cell processes in non-excitable cells, such as migration and invasion, whose derangement promotes the neoplastic progression.

In conclusion, we uncovered in BCa cells a plasma membrane complex whose formation is promoted by β1 integrin-dependent cell adhesion. On ECM adhesion, such complex recruits several ion transport proteins that regulate K^+^, Na^+^ and H^+^ fluxes and cooperate in modulating the downstream signals implicated in cell migration/invasiveness, and thus malignant behavior. As the complex is also found in primary BCa samples, it could be targeted to develop novel therapeutic strategies for one of the most difficult-to-treat cancers, i.e. TNBCa.

## MATERIALS AND METHODS

### Cell culture and transfection

The MDA-MB-231, MDA-MB-468, MCF-7 and SKBR3 cells were obtained from the American Type Culture Collection and were cultured in Dulbecco’s modified eagle medium (DMEM) with 2% L-glutamine and foetal bovine serum (5% for the MDAs cell lines, 10% for MCF-7 and SKBR3). MDA-MB-231 cells stably transfected with hERG1 were cultured in Dulbecco’s modified eagle medium (DMEM) with 2% L-glutamine, 5% foetal calf serum. EBNA cells stably transfected with adult Na_V_1.5 (aNa_V_1.5) and nNa_V_1.5 were cultured in DMEM (Euroclone, Milan, Italy) with 2% L-Glutamine, 10% fetal calf serum. MDA-MB-231 cells stably transfected with hERG1 (MDA-MB-231-T) and with the empty vector were cultured in DMEM with 2% L-glutamine (Euroclone), 5% fetal calf serum (Euroclone, Milan, Italy) and 2 mg/ml geneticin (G418) (Thermo Fisher Scientific, Waltham, MA). Cells were routinely cultured at 37 °C with 5% CO2 in a humidified atmosphere. We certify that all the cell lines used in the present study were routinely screened for mycoplasma contamination, and only mycoplasma-negative cells were used. All cell lines have been acquired from ATCC with proper state authentication (STR profiling).

### Gene silencing experiments

For gene silencing experiments, cells were cultured as above in 60mm cell culture plates (Corning Costar; Corning, NY, USA). Twenty-four hours after plating cells were transfected with two different KCNH2-siRNAs (44858 anti-hERG1 siRNA1 and 44762 anti-hERG1 siRNA2; Ambion; Austin, TX, USA); β1 integrin siRNA (109878 ITGB1; Invitrogen, Waltham, USA) and custom made SCN5A-siRNAs (siNESO1 and siNESO2). A control transfection was also carried out using scrambled siRNAs. The final concentration of the siRNAs was 100 nM in Lipofectamine 2000 reagent (Invitrogen; Carlsbad CA, USA). Transfection followed the manufacturer’s instructions. After 5 hours, the medium was changed and, after overnight incubation, 1 ml Optimem (Gibco; Carlsbad CA, USA) was added. Selection and further cell culture were performed in complete culture medium supplemented with 2.0 mg/ml geneticin (G418). Twenty-four hours later, the supernatant and cells were collected for RNA extraction (see below). 48 hours after transfection, cells were collected, and RNA processed. The siRNA-transfected cells are referred to as “MDA-MB-231-T”. A control transfection was also carried out using scrambled siRNA. The latter are referred to as “MDA-MB-231” since they were identical to MDA-MB-231 un-transfected cells.

### RNA extraction, reverse transcription and real-time quantitative PCR

RNA extraction and reverse transcription were performed as previously described^73,74^. It involved the following primers: neonatal-*SCN5A (nSCN5A) -* 5-CTGCACGCGTTCACTTTCCT-3’ (forward) / 5’-GACAAATTGCCTAGTTTTATATTT-3’ (reverse); adult-*SCN5A* (*aSCN5A*) - 5′-CTGCACGCGTTCACTTTCCT-3′ (forward) / 5′-CAGCCAGCTTCTTCACAGACT-3′ (reverse); *KCNH2 -* 5’-GAGTACAGCCGCTGGATG-3’ (forward) / 5’-ACGTCTCTCCCAACACCA-3’ (reverse). The SYBR Green fluorescent dye (Power SYBR Green PCR Master Mix, Applied Biosystems; Waltham, MA, USA) protocol was used. Glyceraldehyde-3-phosphate dehydrogenase gene was used as a standard reference: *GADPH*: (forward) 5’-ACGACCAAATCCGTTGACTC-3’ / (reverse) 5’-GCTGTCTGCTCCTCCTGTTC-3’^75,76^. Calibration was performed using HEK-293 and HEK-hERG1 cell lines for *KCNH2* and EBNA-nNa_V_1.5 for nNa_V_1.5. ΔΔCt method was used for the relative quantification of the expression levels. Amplification of a specific products or primer–dimer artefacts was excluded by melting curve analysis. Each reaction was performed in triplicate.

### Preparation of cells for experiments

For all experiments, excluding Matrigel invasion assay, all the cell types were starved overnight by culturing them in serum-free-BSA medium, containing 250 mg/ml of heat inactivated (HI) BSA (Fraction V, Euroclone, Milan, Italy). The day after, cells were harvested by detaching them with 5 mM EDTA in phosphate-buffered saline (PBS) (Euroclone, Milan, Italy) and resuspended in serum-free medium. Next, cells were seeded on FN-coated dishes/slides. To this purpose culture dishes/glass slides were first coated with fibronectin (Sigma-Aldrich, Darmstadt, Germany, human plasma) at 5 µg/cm^2^ concentration diluted in sterile PBS. The dishes were left air-drying for 1 h at room temperature before introducing cells and medium. Hence, cells detached and resuspended in serum-free-BSA medium, were added at different concentrations depending on the type of experiment to be performed. On the day of the experiment, cells were properly counted to obtain the exact concentration needed as in Dimitri and Duranti et al ^77^. For experiments with silenced cells overnight starvation was not carried out.

### Treatments

All treatments have been performed diluting stock solutions in fresh medium on the day of the experiment. We used the following inhibitors: 40 µM E4031, (starting solution 10 mM in sterile double distilled water; Tocris, Bristol, UK; hERG1 blocker^37^); 20μM tetrodotoxin (TTX) (starting solution 3 mM in sterile double distilled water; Tocris, Bristol, UK; nNa_V_1.5 blocker^78^), 20µg/ml scDb-hERG1-β1 (starting solution 2 mg/ml in PBS; MCK Therapeutics Srl, Pistoia, Italy; hERG1/β1 integrin complex blocker^10,22^) and 5 μM cariporide (starting solution 5 mM in DMSO; Tocris, Bristol, UK; specific NHE1 inhibitor^79^).

### Protein extraction, co-immunoprecipitation and Western Blotting

Cells were starved O/N in culture media plus heat-inactivated BSA (250 mg/ml; Fraction V). Cells were harvested by detaching with 5 mM EDTA in phosphate-buffered saline (PBS) and resuspended in culture media plus heat-inactivated BSA were seeded onto dishes pre-coated with 5 µg/cm^2^ fibronectin (in sterile PBS). For (co)-immunoprecipitation experiments, cells were seeded and incubated on fibronectin for 120 minutes. Protein extraction, quantification and total lysate incubation with protein A/G agarose beads (Santa Cruz Biotechnology; Dallas, TX, USA) were performed as reported by Becchetti et al^15^. Protein extraction, quantification and total lysate incubation with protein A/G agarose beads (Santa Cruz Biotechnology; Dallas, TX, USA) were performed as reported by Becchetti et al^15^. The composition of the lysis buffer was the following: NP40 (150 mM), NaCl (150 mM), Tris-HCl pH 8 (50 mM), EDTA pH 8 (5 mM), NaF (10 mM), Na_4_P_2_O_7_ (10 mM), Na_3_VO_4_ (0.4 mM), and a protease inhibitor cocktail (complete Mini, Roche; Basel, Switzerland)^80^. The TS2/16 antibody (details are reported in Table S1) was used to immunoprecipitate β1-integrin; the mAb-hERG1 antibody (details are reported in Table S1) was used to immunoprecipitate hERG1 protein. The Clone TS2/16 antibody was left incubating overnight and after that the immuno-complex was captured by adding 50μl of protein A/G agarose (Merck Sigma, Burlington, MA) beads for 2 hours at 4°C (with rolling agitation). The incubation was followed by two washing steps: 3 times with ice-cold wash buffer and 3 times with ice-cold PBS. Then, 10μl of 2X Laemmli Buffer were added and the proteins were boiled for 5 min at 95°C. SDS-PAGE was performed and following electrophoresis, proteins were transferred onto PVDF membrane (previously activated) in blotting buffer at 4°C for 1 h at 150 V. 5% BSA in T-PBS (0.1% tween, Merck Sigma, Burlington, MA) solution was then used to block the PVDF membrane, for 3 hours at room temperature (this step is required to cover the unspecific antibody binding sites on the membrane). Blots were then incubated overnight at 4°C with primary antibodies: C54^81^, RM-12^82^, mAb-nNa_V_1.5 (Duranti et al, personal communication, January 2025), anti-NHE1^83^ (type and antibodies dilution are described in Table S1). The following day the membrane was washed with T-PBS (0.1% tween) (15 min x 3 times) and appropriate secondary antibody: (i) anti-rabbit antibody conjugated with peroxidase enzyme (Merck Sigma, Burlington, MA) was dissolved in 5% BSA in T-PBS (0.1% tween) (dilution 1:10.000) for at least 45 min and washed (15 min x 3 times), revealing was performed using ECL solution (Amersham, Buckinghamshire, UK) for anti-C54 primary antibody^81^ and (ii) for all other primary antibodies, IRDYe 800 CW anti-mouse or anti-rabbit antibody (LI-COR Biosciences, Nebraska, USA) was dissolved in 5% BSA in T-PBS (0.1% tween) (dilution 1:20.000) for at least 45 min and washed (15 min x 3 times) before membrane scanning using LI-COR Odyssey (Biosciences, Nebraska, USA).

### Densitometric analysis

All WB quantification were performed by dividing protein band by tubulin signals. Co-immunoprecipitated hERG1, β1 integrin, NHE1 and nNa_V_1.5 proteins, alone and in combination, were quantified by optical densitometry. The co-immunoprecipitated signal (i.e. hERG1) was first separated by the signal of the protein used for the immunoprecipitation process (i.e. β1 integrin) and then normalized to the signal of the corresponding protein obtained in the total lysate (i.e. hERG1). ImageJ was used to analyze data and graphs were plotted using OriginPro8 software (OriginLab Corporation; Northampton, MA, USA). The expression levels of the proteins in WB and co-IP have been quantified as reported above. Then they have been normalized as following: (WB quantification/IF quantification)*100 where WB is referred to total lysates of hERG1 normalized on tubulin; IF is referred to hERG1 protein expression on plasma membrane.

### Immunofluorescence

24-well plates were prepared with one coverslip pre-coated with Fibronectin 5μg/cm^2^ placed in each well and incubated at least 1 hour. Then the coating solution was removed and 2×10^4^cells were seeded on each coverslip and left to settle for 2h. Cells were then fixed with 500μL 4% paraformaldehyde (PFA) (Thermo scientific, Rockford, IL, USA) solution in PBS 1X, pH 7.4 at room temperature. After 10 minutes, the PFA solution was removed and each well was washed with PBS 3 times (5 minutes each). Cells were then ‘blocked’ by washing once with 500μl of 10% BSA solution in PBS (pH 7.4), for 1 hour at room temperature. A solution of the primary antibody Alexa-488 or Alexa-647 conjugated (at final concentration indicated in Table S1) was made up and 100μl were added onto each coverslip placed on a podium in a humidity chamber and cells were allowed to incubate overnight in a cold room (6-7°C) for hERG1 and nNav1.5 staining and 2 hours at room temperature for hERG1/β1 integrin complex labelling. For paxillin and Arp2/3 detection, overnight incubation in a cold room (6-7°C) was carried and anti-mouse/rabbit secondary antibody Alexa-488 conjugated (Thermo Fisher Scientific, Waltham, MA) incubation was performed (1 hour, room temperature, 1.500 final dilution). Then, each coverslip was washed with PBS 3 times each for 5 minutes and nuclei were stained with Hoechst (Merck Sigma, Burlington, MA; 15 min, room temperature, 1:1000 final dilution) and slides were mounted using Prolong Diamond antifade mountant (Invitrogen, Waltham, Massachusets, USA). To investigate cytoskeletal actin architecture using confocal microscopy, cells were fixed using 4% PFA, followed by permeabilization (0.1% Triton-X; Merck Sigma, Burlington, MA), blocking with 1% BSA and staining with rhodamine-conjugated phalloidin following manufacturer’s instructions (Invitrogen, Waltham, MA, USA) and Hoechst (15 min, room temperature, 1:1000 final dilution) and slides were mounted using Prolong Diamond antifade mountant (Invitrogen, Waltham, Massachusets, USA). Pictures were taken on a confocal microscope (Nikon TE2000, Nikon; Minato, Tokyo, Japan).

### Immunofluorescence signal quantification

In all immunofluorescence experiments, the fluorescence relative to 20 cells (in 10 different fields and 3 different experiments) per each condition was determined using ImageJ Software. For total IF signal, we measured the mean fluorescence intensity for each cell. When needed, the fluorescent intensity at the plasma membrane level was quantified considering exclusively the signals arising at the periphery of the cells. To this purpose we draw a mask (highlighted by white circular lines in the figures) that was performed determined using the “freeform line profile” function drawn around the cell surface that enucleates only the fluorescent signal related to the membrane which was subsequently quantified using ImageJ Software^10^. Colocalization was quantified using Manders’ Overlap coefficient, which was measured using the Co-loc plugin (Fiji Software) for each cell analyzed. For total IF signal, we measured the fluorescence intensity for each cell, dividing it by the cell area, to obtain a final mean fluorescence (± standard error mean). Each single experiment replicate has been performed using the same conjugated antibody and acquiring images during the same confocal microscopy session. Moreover, for each acquisition the same script has been maintained. To analyze filopodia, actin filaments within 2 μm of the cell border were taken into account. The (linear) density of filopodia was calculated as the total number of filopodia normalized by the cell perimeter length. The cortactin staining was quantified as the “Invadopodia Index” (InvI), i.e. percentage of cells showing clear brightly stained spots. 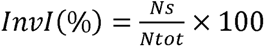

Ns = cells showing invadopodia spots

Ntot = total number of cells.

### Cell proliferation

The 3-(4,5-dimethylthiazo1-2-yl)-2,5-diphenyltetrazolium bromide (MTT) (Merck Sigma, Burlington, MA) reduction assay was used to quantify cell proliferation. The assay enables determination of the level of metabolic activity in eukaryotic cells via cytoplasmic and mitochondrial reduction of MTT (yellow) into formazan crystals (blue). Cells were plated in 24 well plates at a density of 2×10^4^ cells/well and let them settle overnight. The DMEM solution was then replaced with 1mL of treatment solution for 24 hours. Cells were then treated with 100μl of MTT and 400μl of DMEM and incubated (in darkness) for 3 hours as normal at 37°C. The MTT solution was replaced with 50μl of DMSO (Merck Sigma, Burlington, MA). Then, the plates were placed on a rocker in darkness for 10 minutes. The absorbance of each well was read at 570 nm on a multi-well plate reader. Cell number was deduced from a pre-determined linear calibration curve. The latter was generated by serial dilution of cells (50, 25, 12.5, 6.250, 3.125 and 1.5×10^4^) left to settle for 4 hours before applying the MTT protocol. Each calibration and each drug treatment were repeated at least 3 times.

### Lateral motility

Lateral motility was determined using 35 mm dishes and drawing 15 horizontal lines and 3 perpendicular lines on the dish bottom, as in Iorio et al^11^. Plates were coated with FN (Sigma-Aldrich, Darmstadt, Germany, human plasma) and 5×10^5^ cells were seeded and allowed to attach 5-10 min. Then, a manual scratch was done and the width of the wound was determined (W0). At this time, the different treatments were added. Then, dishes were incubated for further for 120 min. At the end of incubation, the width of the wounds (Wt) was determined on unfixed cells. We took care to do these measurements within non more than 30 min overall. Motility Index (MI) was assessed using the following formula: MI = 1 – Wt/W0, where Wt is the width of the wounds.

### Matrigel invasion

Following 24h pre-treatment, cells were plated in 24-well plates containing Transwell^®^ filters (Beckton Dickinson; Oxford, UK) with 8μm pores. Filters were coated a day earlier with 50μL Matrigel^®^ (Beckton Dickinson; Oxford, UK) diluted with FBS-free DMEM to 1.25 mg/mL from a 10 mg/mL stock and left to solidify overnight. Inserts were then hydrated with 0.5mL FBS-free DMEM for 2-3 hours at 37°C. Cells were plated onto the coated filters at a density of 2×10^4^ cells/insert; a chemotactic gradient was produced by adding 300μL of 5% FBS supplemented solutions in the lower chamber and 300μL of 1% FBS supplemented solution in the upper chamber. Treatment with E4031 or TTX was continued by adding the agent to the upper chamber and cells were allowed to invade for 18 hours. All the solutions were removed and the upper parts of the inserts were swabbed to remove the non-invaded cells. Invaded cells were fixed in 300μL ice-cold methanol (100%) for 15 minutes. Staining was with 200μL of 0.5 g/ml crystal violet in 25% methanol for 15 minutes at room temperature. Inserts were washed with distilled water and dried; stained cells were counted in at least 12 random fields of view/insert at 40x magnification on an inverted microscope. The number of cells counted in each field of view was converted to “cells per mm^2^” by multiplying the number of counted cells by a factor of 4.07. For all experiments, a background level of invasion was calculated, seeding the cells without chemotactic gradient, and a basal level of invasion was then subtracted from the measurements.

### Electrophysiology

On the experimental day, MDA-MB-231-T and WT cells were detached, resuspended in DMEM 5% FBS with and without 2 mg/ml G418, respectively, and seeded on 35 mm Petri dishes. Cells were identified at 40× magnification with a Nikon Eclipse TE300 microscope (Nikon Instruments Inc., Amstelveen, The Netherlands), equipped with a Photometrics CoolSNAP CF camera (Teledyne Photometrics, Tucson, AZ, USA). Electrophysiological recordings were performed at room temperature (∼25 °C) in the whole-cell configuration of the patch clamp technique, after 2 h during which cells were maintained in an incubator at 37 °C, 5% CO_2_. The patch pipettes were pulled from borosilicate glass capillary tubes to a resistance of 3.5–5 MΩ. Capacitances were manually compensated after reaching a stable gigaseal. For stimulation and data acquisition/analysis we used Multiclamp 700A or Multiclamp 1D amplifiers equipped with pCLAMP 9.2/Digidata hardware and software (Molecular Devices, Sunnyvale, CA, USA). hERG1 currents were low-pass filtered at 2 kHz and sampled at 25 kHz. Holding membrane potentials was −80 mV. hERG1 inward tail currents were elicited at −120 mV, preceded by conditioning voltage steps ranging from 0 mV to −100 mV (10 mV step increment; 5 s step duration; 15 s intersweep intervals). Pipette contained (mM): 130 K^+^ aspartate, 10 NaCl, 4 CaCl_2_, 2 MgCl_2_, 10 Hepes, 10 EGTA, pH 7.3. The external solution contained (in mM): 130 NaCl, 5 KCl, 2 CaCl_2_, 2 MgCl_2_, 10 HEPES, 5 glucose (E_K_ = −80 mV), pH 7.4. The hERG1 current amplitude, normalized to the maximum current amplitude, was used to construct the activation curve (**Supplementary Figure S5D**). For Nav1.5 current recording, the external solution contained (in mM): 144 NaCl, 5.4 KCl, 1 MgCl_2_, 2.5 CaCl_2_, 5 HEPES, and 5.6 glucose, pH 7.3. The pipette solution contained (mM): 145 CsCl, 5 NaCl, 2 MgCl2, 11 EGTA, and 10 HEPES, pH of 7.4. Signals were low-pass filtered at 5 kHz and sampled at 10 kHz. Nav1.5 currents were elicited by applying voltage steps from −100 mV to +45 mV (5 mV step increment; 400ms duration) from a holding potential of −100 mV. Intersweep interval was 2 s.

### Calcium imaging

We followed the procedure reported in Scarpellino et al^84^. In brief, cells were grown on glass fibronectin- or BSA-coated coverslips, after overnight starvation, at a density of 5000 cells/cm^2^ for 2 h. Next, cells were loaded (40 min at 37°C) with 2 µM Fura-2AM (Invitrogen; Carlsbad, CA, USA), for ratiometric cytosolic Ca^2+^ [Ca^2+^]c measurements. A Nikon Eclipse TE-2000S (Nikon; Minato, Tokyo, Japan) inverted microscope was used to acquire fluorescence. [Ca^2+^]c was expressed as the ratio between emitted fluorescence at 510 nm after excitation at 340 nm and 380 nm. Image acquisition (3 s frequency) was performed using Metafluor Imaging System (Molecular Devices; Sunnyvale, CA, USA). For each experiment, several regions of interest (ROIs) have been selected corresponding to single cells in a chosen image field. Real time background subtraction was applied in order to limit noise. Calcium imaging analysis and quantification (peak amplitude, area, rise time, decay time) were performed using Clampfit software (pCLAMP 9.2, Molecular Devices) and analyzed with GraphPad Prism 8 (GraphPad Software Inc., La Jolla, CA, USA). Area underlying calcium influx during the sustained phase in the presence of extracellular Ca^2+^ was evaluated at 200 s after the onset of the response. Total area under calcium spikes in 0 Caout was measured by using the Event Detection protocol in Clampfit software.

### Measurement of pH_i_

To determine pH_i_, we used 2’,7’-Bis (2-carboxyethyl)-5 (6)- carboxyfluorescein acetoxymethyl ester (BCECF-AM) (Sigma-Aldrich, Darmstadt, Germany). Cells were starved overnight. After starvation, 50.000 cells were seeded in no-serum medium onto BSA or FN coated surfaces of 96-well plates (clear bottom 96-well plate, polystyrene, TC-treated, clear flat bottom wells, sterile, w/lid, black; Corning, New York, USA) and incubated at 37°C and 5% CO_2_ for 2 h, in the absence or presence of different treatments. At 2 h, the medium was removed and BCECF-AM (1μM final concentration in loading solution (HBSS 1X (Euroclone, Milan, Italy) plus 0.01% NaHCO_3,_ pH 7.4)) was added for 30 min at 37°C and 5% CO_2_. After incubation, cells were washed twice with loading solution at room temperature. The time zero pH_i_ (T_0_) was measured in detached cells, before seeding, at room temperature. The pH_i_ of detached cells was determined in 96-well plate with microplate reader and the appropriate standard curve. Fluorescence intensity was immediately measured with a microplate reader at the following wavelengths: 440–490 nm for excitation and 535 nm for emission. A calibration curve was set up using a high K^+^/Nigericin solution (135 mM KCl, 2 mM K_2_HPO_4_, 20 mM HEPES, 1.2 mM CaCl_2_ and 0.8 mM MgSO_4_), in a range of pH from 5.0 to 8.5. All pH values were calculated using 490/440 nm fluorescence ratio and applying standard curve and linear equations.

### Statistical analysis

Unless otherwise indicated, data are given as mean values ± SEM, with n indicating the number of independent experiments. Statistical comparisons were performed with OriginPro 2015 and SAS 9.2 (SAS Institute) software. The normality of data distribution was checked with K-S test. In the case of normal distributions, each data set was first checked for variance homogeneity, using the F test for equality of two variances and the Brown-Forsythe test for multiple comparisons. For comparisons between two groups of data, we used the Student’s t test. For data with unequal variances, the Welch correction was applied. For multiple comparisons, one-way ANOVA followed by Bonferroni’s post hoc test was performed to derive P values. As reported in the figure legends, in case of unequal variances, ANOVA was followed by the Hochberg’s GT2 post hoc method. In the case of nonnormal distributions, nonparametric Kruskal-Wallis ANOVA followed by DSCF’s post hoc method was applied. The relevant P values are reported in the figure panels and legends. The Pearson coefficient (R) was calculated to evaluate relationships between continuous variables (R□=□−□1 negative relationship; R□=□0 no relationship; R□=□1 positive relationship).

## ACKNOWLEDGMENTS

The authors thank Dr Matteo Lulli for confocal microscopy, (Department of Experimental and Clinical Biochemical Sciences, Section of General Pathology, University of Florence, Florence, Italy); Prof Luca Munaron and Prof Alessandra Fiorio Pla for help in experiments measuring intracellular Ca^2+^ (Department of Life Sciences and Systems Biology, University of Torino, via Accademia Albertina 13, 10123 Torino, Italy).

