## Supplementary data for "An integrin centered complex coordinates ion transport and pH to regulate f-actin organization and cell migration in breast cancer"

| Target | Antibody type / supplier | Dilution |
| --- | --- | --- |
| nNav1.5 | Monoclonal / Celex Holdings Limited (12941723) | WB: 1:200  IF: 1:200 (Alexa-488, Alexa-647 conjugated)  IHC: 1:100 |
| hERG1 | Polyclonal / C54, DI.V.A.L Toscana srl, Sesto Fiorentino, Italy | WB: 1:1000 |
| hERG1 | Monoclonal / mAb-hERG1, MCK Therapeutics, Pistoia, Italy | IP: 5 µg for 1mg of protein  IF: 1:1000 (Alexa-488 conjugated)  IHC: 1:500 |
| β1 Integrin | Monoclonal / LEAF™ Purified anti-human CD29, Clone TS2/16, Biolegend, San Diego, CA, USA | WB: 5μg for 1mg of protein |
| β1 Integrin | Polyclonal / RM-12, Immunological Science, Rome, Italy | WB: 1:1000 |
| Paxillin | Monoclonal/anti-paxillin, BD Transduction Laboratories, USA | IF: 1:200; WB: 1:1000 |
| Phalloidin | Rhodamine-conjugated phalloidin, Invitrogen, Waltham, MA, USA | IF: 1:10 |
| Cortactin | Polyclonal / Cortactin Polyclonal Antibody, Invitrogen, Waltham, MA, USA | IF: 1:500 |
| Actin | Monoclonal / Actin Monoclonal Antibody (mAbGEa), Invitrogen, Waltham, MA, USA | WB: 1:3000 |
| Arp2/3 | Polyclonal / Arp2 polyclonal antibody PA5-118867; Arp3 polyclonal antibody PA5-17311, Invitrogen, Waltham, USA | IF: 1µg/ml; 1:100 |
| NHE1 | Polyclonal / anti-NHE1, Novus Biologicals, Centennial, CO, USA | WB: 1:500  IHC: 1:400 |
| Vinculin | Monoclonal / anti-vinculin (7F9), Santa Cruz Biotechnology, Dallas, TX, USA | WB: 1:2000 |
| α-actinin 1 | Monoclonal / anti- α-actinin (H-2), Santa Cruz Biotechnology, Dallas, TX, USA | WB: 1:500 |
| Tubulin | Monoclonal / anti-tubulin (F-1), Santa Cruz Biotechnology, Dallas, TX, USA | WB: 1:500 |
| scDb-hERG1/β1 | Single chain diabody, MCK Therapeutics Srl, Pistoia, Italy | Cell Treatment: 20µg/ml  IF: 20µg/ml (Alexa-488 conjugated)  IHC: 20µg/ml |
| pAKT1/2/3 | Monoclonal / anti-pAKT (H-136) (cat. Sc-8312) / Santa Cruz Biotechnology, Dallas, TX, USA. | WB: 1:500  IF: 1:200 |
| AKT1/2/3 | Monoclonal / anti-AKT (Thr 308) (cat. sc-271966) / Santa Cruz Biotechnology, Dallas, TX, USA. | WB: 1:500 |

**Table S1** Antibodies used in the experimental procedures reported in the present study.

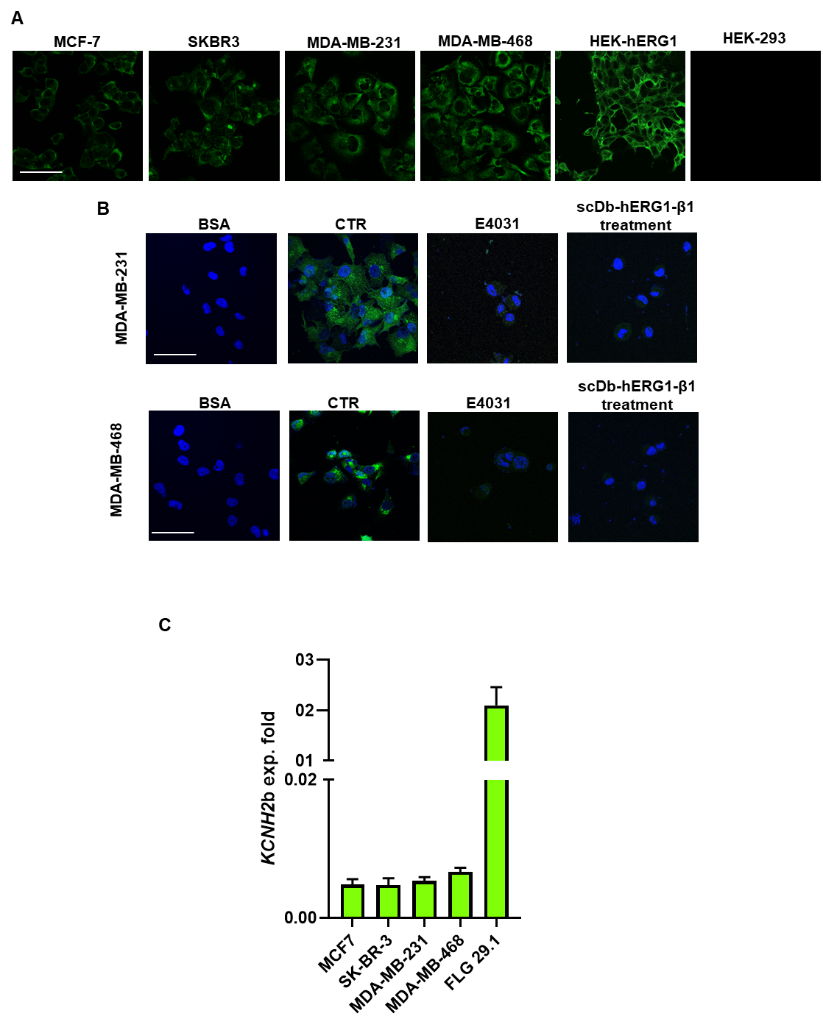

**Supplementary Figure S1. A)** Representative images of expression levels of hERG1, determined by Immunofluorescence (IF) in breast cancer cell lines (MCF-7, SKBR3, MDA-MB-231, MDA-MB-468). Cells were seeded onto FN for 2 hours. Cells were stained for hERG1 (in green) with mAb-hERG1 primary antibody conjugated with AlexaFluor-488. **B)** hERG1/β1 complex formation evaluated by staining with fluorescent scDb–hERG1–β1 in MDA-MB-231 and MDA-MB-468 cells seeded on FN and BSA as negative control. Representative IF images at T_120_ (left panel) (scale bars: 100 μm) and IF densitometric analysis (right panel) are reported. Representative images (scale bar 100 µm) and quantification are reported. For each condition the fluorescence relative to 20 cells was analysed. Pictures were taken on a confocal microscope (Nikon TE2000, Nikon; Minato, Tokyo, Japan). ImageJ software was used to analyse the images. **C)** Expression levels of the *KCNH2b* transcripts, determined by Real Time PCR in different breast cancer cell lines (MCF-7, SKBR3, MDA-MB-231, MDA-MB-468), FLG 29.1. FLG 29.1 were used as positive control for *KCNH2b*. Cells were seeded onto Fibronectin (FN) for 2 hours. Data, reported as 2-DCt, are mean values ± s.e.m. (n=3).

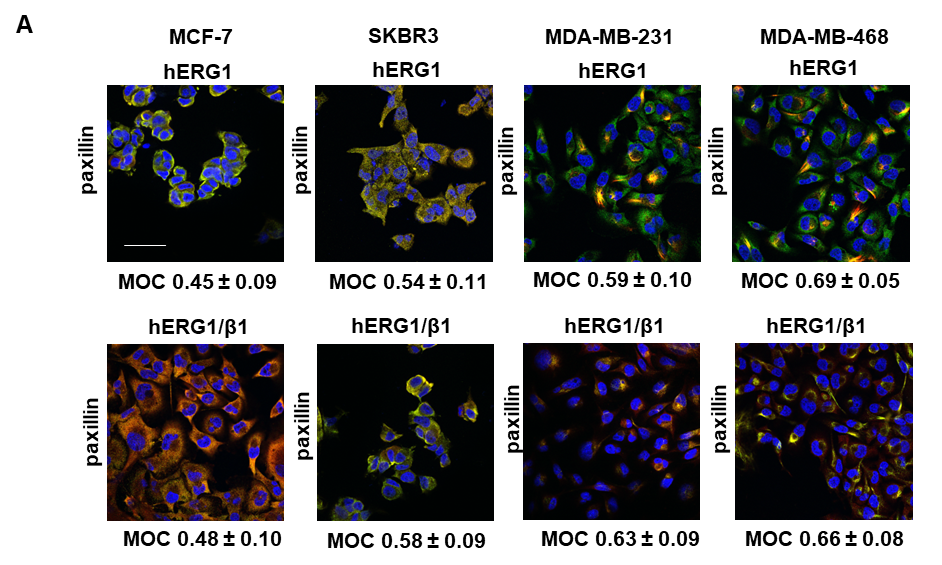

**Supplementary Figure S2. A)** Representative IF images (scale bar: 100 μm) of the co-localization of hERG1 (in green probed with mAb-hERG1 conjugated with AlexaFluor-488) and paxillin (in red probed with mAb-paxillin and secondary Ab conjugated with AlexaFluor-546) (top panel), and hERG1/β1 (in green probed with scDb-hERG1/β1 conjugated with AlexaFluor-488) and paxillin (in red probed with mAb-paxillin and secondary Ab conjugated with AlexaFluor-546) (bottom panel) in breast cancer cell lines (MCF-7, SKBR3, MDA-MB-231, MDA-MB-468) seeded onto FN for 2 hours. MOC reporting hERG1/paxillin and hERG1/β1/paxillin correlations is showed next to each image. For each condition the fluorescence relative to 20 cells was analysed. MOC = Manders’ Overlapping Coefficient.

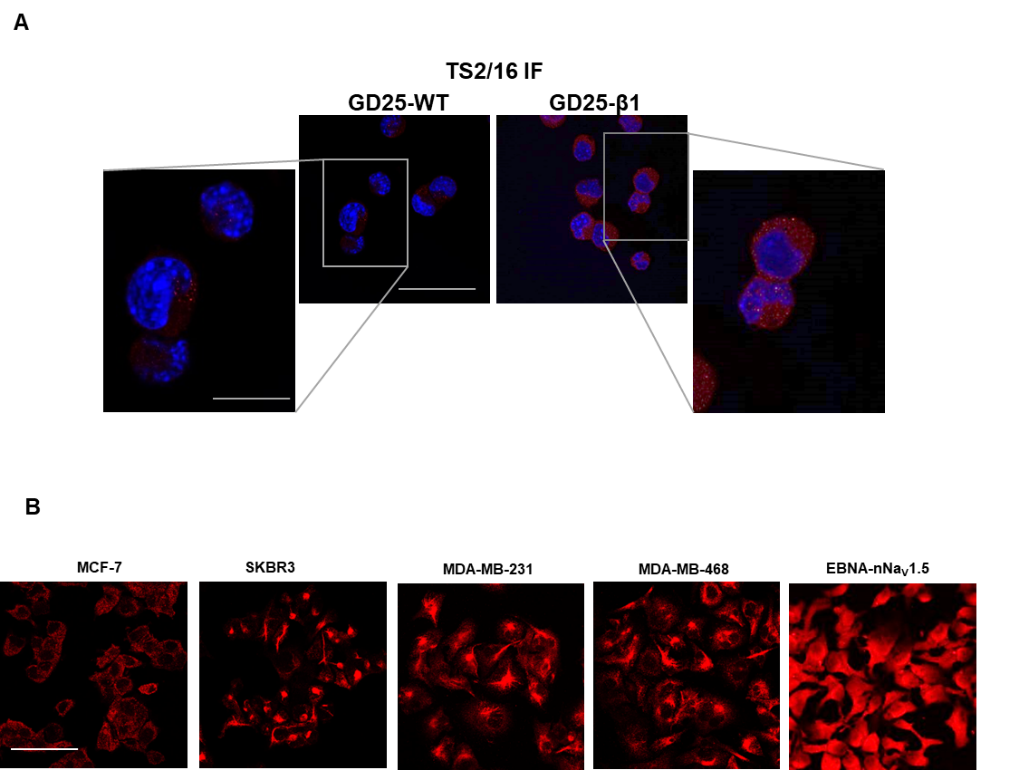

**Supplementary Figure S3. A)** Representative images of IF experiments on GD25-WT and GD25-β1 cells, showing the specificity of TS2/16 antibody. **B)** Representative images of successful silencing of nNa_V_1.5 on all BCa cell lines. **C)** Representative images of expression levels of nNa_V_1.5, determined by Immunofluorescence (IF) in breast cancer cell lines (MCF-7, SKBR3, MDA-MB-231, MDA-MB-468). Cells were seeded onto FN for 2 hours. Cells were stained (in red) with mAb-nNa_V_1.5 primary Ab conjugated with AlexaFluor-647. Representative images (scale bar 100 µm) and quantification are reported. The quantification was performed considering only the membrane signal, highlighted by the white masks shown in the pictures. For each condition the fluorescence relative to 20 cells was analysed. Pictures were taken on a confocal microscope (Nikon TE2000, Nikon; Minato, Tokyo, Japan). ImageJ software was used to analyse the images.

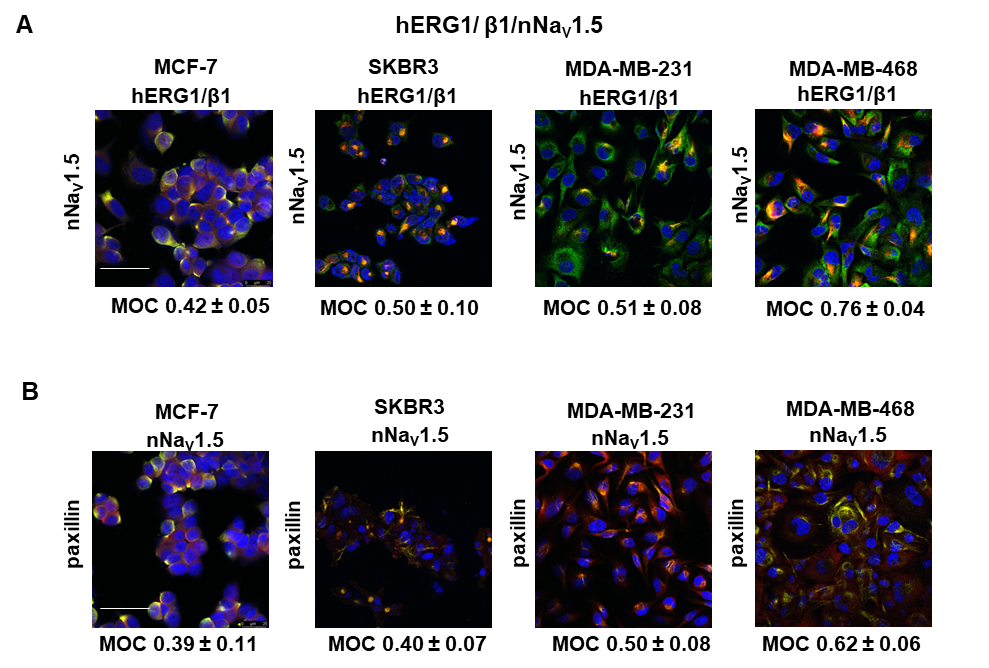

**Supplementary Figure S4. A)** Representative IF images (scale bar: 100 μm) of the co-localization of hERG1, β1 integrin (in green probed with scDb-hERG1-β1 conjugated with alexa-488) and nNa_V_1.5 (in red probed with mAb-nNa_V_1.5 conjugated with AlexaFluor-647) in breast cancer cell lines (MCF-7, SKBR3, MDA-MB-231, MDA-MB-468) seeded onto FN for 2 hours. Mander’s Overlapping Coefficient (MOC) reporting the proteins correlation is showed under each image. For each condition the fluorescence relative to 20 cells was analysed. **B) )** Representative IF images (scale bar: 100 μm) of the co-localization of nNa_V_1.5 (in green probed with mAb-nNa_V_1.5 conjugated with AlexaFluor-488) and paxillin (in red probed with mAb-paxillin and secondary Ab conjugated with AlexaFluor-546) (bottom panel) in breast cancer cell lines (MCF-7, SKBR3, MDA-MB-231, MDA-MB-468) seeded onto FN for 2 hours). Representative IF images (scale bar: 100 μm) of the co-localization of nNa_V_1.5 (in green probed with mAb-nNa_V_1.5 conjugated with AlexaFluor-488) and paxillin (in red probed with mAb-paxillin and secondary Ab conjugated with AlexaFluor-546) (bottom panel) in breast cancer cell lines (MCF-7, SKBR3, MDA-MB-231, MDA-MB-468) seeded onto FN for 2 hours. MOC reporting nNa_V_1.5/paxillin correlations is showed underneath each image. For each condition the fluorescence relative to 20 cells was analysed. For each condition the fluorescence relative to 20 cells was analysed.

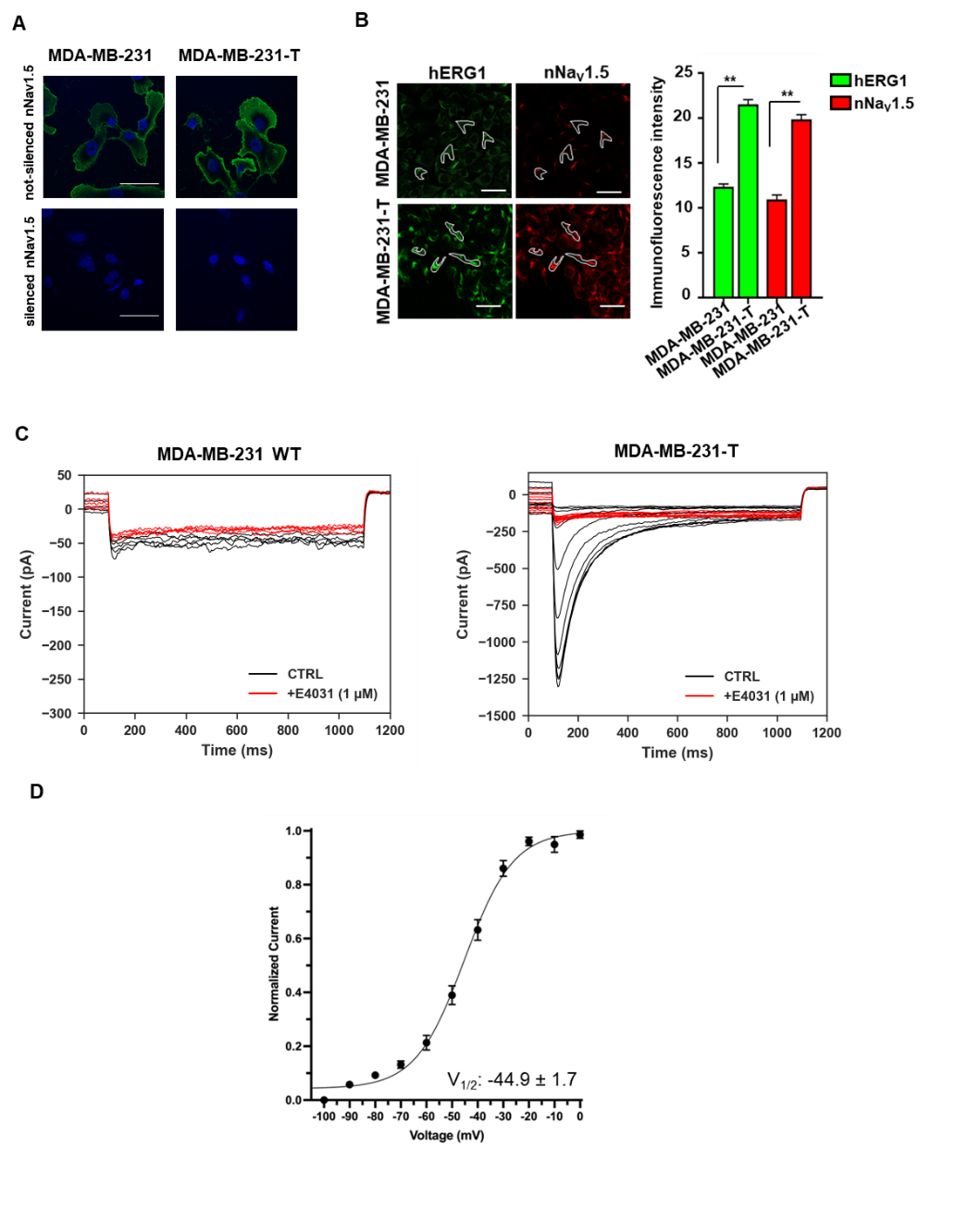

**Supplementary Figure S5. A)** Membrane expression levels of the two proteins, hERG1 and nNa_V_1.5, determined by IF in MDA-MB-231-T cell lines. Cells were stained for hERG1 (in green) with mAb-hERG1 primary antibody conjugated with AlexaFluor-488 and nNa_V_1.5 (in red) with mAb-nNa_V_1.5 primary Ab conjugated with AlexaFluor-647. Representative images (scale bar 100 µm) and quantification are reported. The quantification was performed considering only the membrane signal, highlighted by the white masks shown in the pictures. For each condition the fluorescence relative to 20 cells was analysed. Data are mean values ± s.e.m. (n=3). **B)** Expression of hERG1 determined by IF in MDA-MB-231 and MDA-MB-231-T cell lines, after silencing of nNa_V_1.5. Cells were stained for hERG1 (in green) with mAb-hERG1 primary antibody conjugated with AlexaFluor-488. Representative images (scale bar 100 µm). Pictures were taken on a confocal microscope (Nikon TE2000, Nikon; Minato, Tokyo, Japan). ImageJ software was used to analyse the images. **P < 0.01. **C)** Representative traces of hERG1 before (black) and after (red) the application of E4031 (1µM) in MDA-MB-231 cells (n = 7) (left) and MDA-MB-231-T cells (n = 9) (right) and D) mean activation curves of hERG1. Data are mean values ± s.e.m. (n=3).

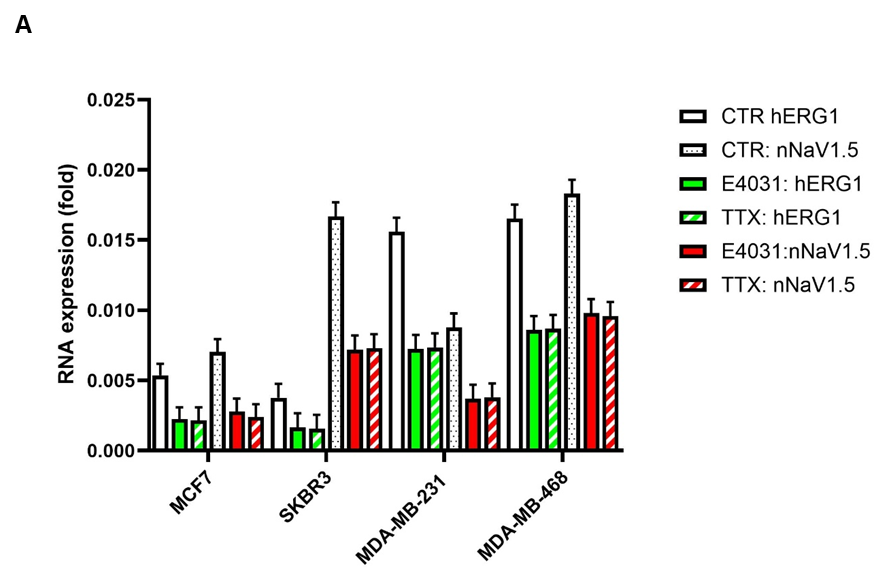

**Supplementary Figure S6.** **A)** Expression levels of the *KCNH2* and *nSCN5A* transcripts after treatment with E4031 and TTX, determined by Real Time PCR in different breast cancer cell lines (MCF-7, SKBR3, MDA-MB-231, MDA-MB-468). Cells were seeded onto Fibronectin (FN) and treated for 2 hours as reported in Materials and Methods. Data, reported as 2-DCt, are mean values ± s.e.m. (n=3).

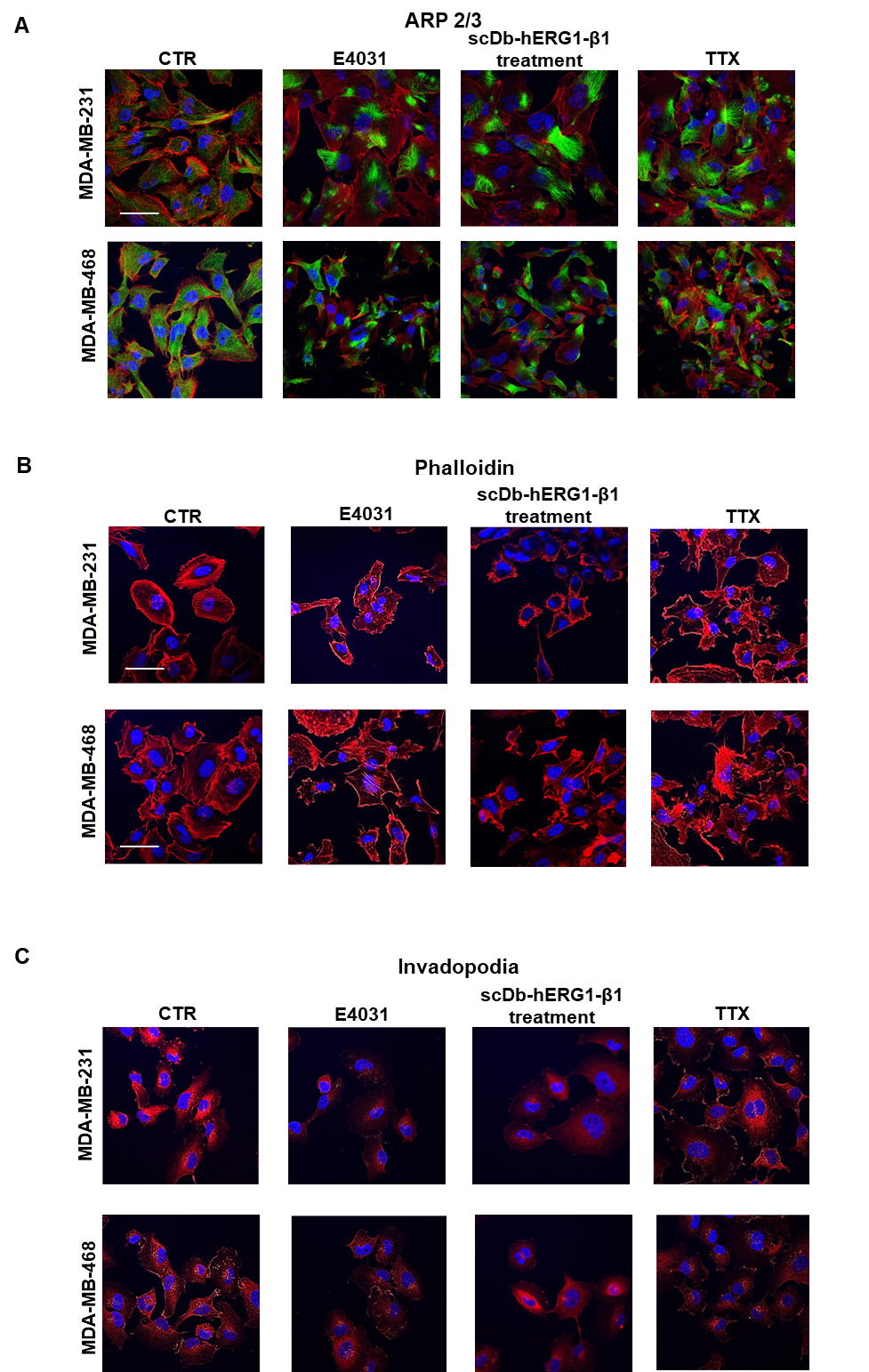

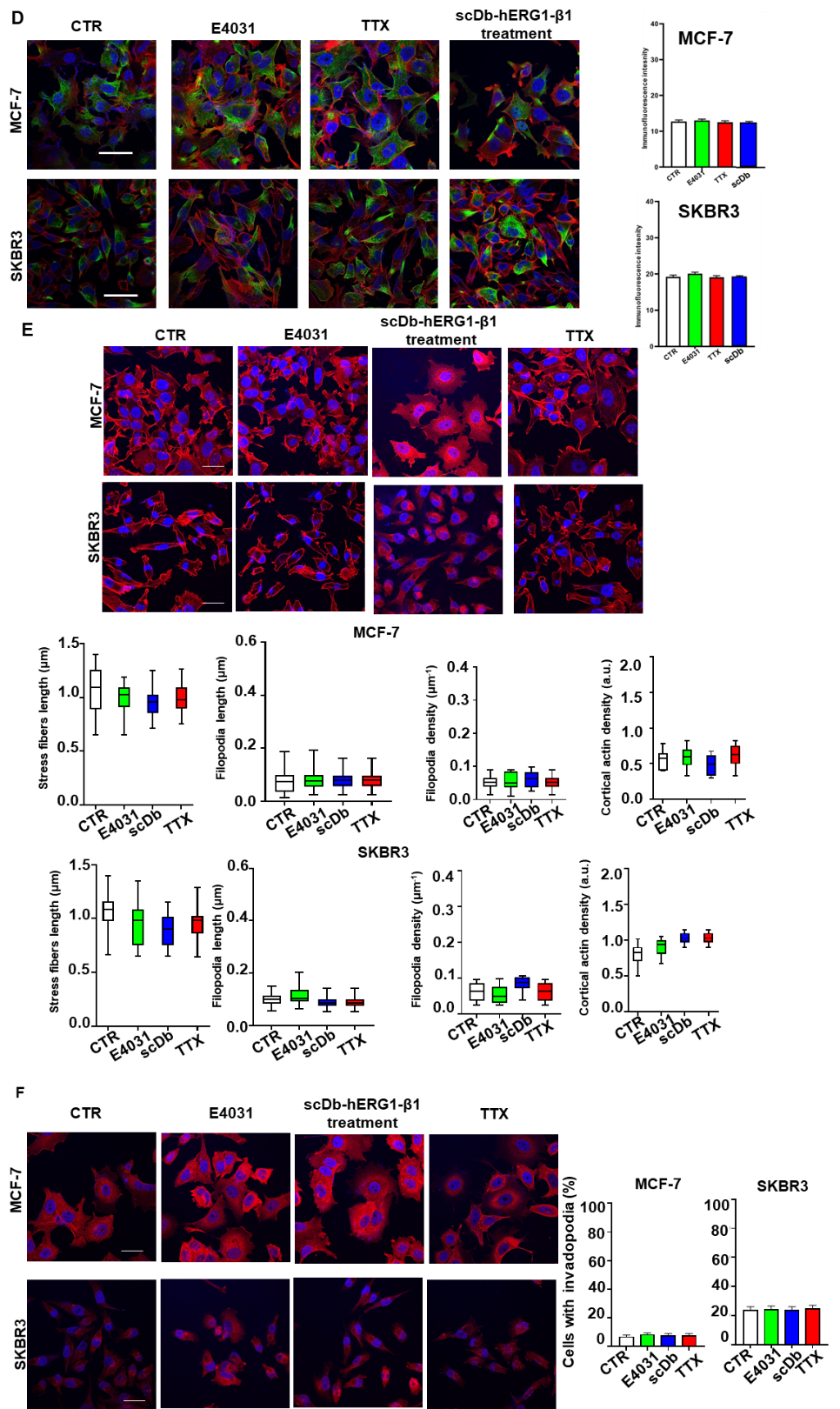

**Supplementary Figure S7. A)** Staining with anti-Arp2/3 (green) antibody with secondary Ab AlexaFluor-488 and CellMask™ Deep Red Actin Tracking Stain (red) of MDA-MB-231 and MDA-MB-468 cells seeded on FN for 2 hours and treated with E4031: 40μM, scDb-hERG1/β1: 20µg/ml and TTX: 20μM. **B)** Phalloidin staining (Ab conjugated with Rhodamine) of MDA-MB-231 and MDA-MB-468 cells seeded on FN for 2 hours and treated with E4031: 40μM, scDb-hERG1/β1: 20µg/ml and TTX: 20μM. Corresponding graphs of actin stress fibres length, filopodia length and cortical actin density. **C)** Staining with anti-cortactin antibody of MDA-MB-231 and MDA-MB-468 cells seeded on FN for 2 hours and treated with E4031: 40μM, scDb-hERG1/β1: 20µg/ml and TTX: 20μM. **D)** Staining with anti-Arp2/3 (green) antibody with secondary Ab AlexaFluor-488 and CellMask™ Deep Red Actin Tracking Stain (red) of MCF-7 and SKBR3 cells seeded on FN for 2 hours and treated with E4031: 40μM, scDb-hERG1/β1: 20µg/ml and TTX: 20μM. Corresponding Arp2/3 graphs on the bottom panel. **E)** Phalloidin staining (Ab conjugated with Rhodamine) of MCF-7 and SKBR3 cells seeded on FN for 2 hours and treated with E4031: 40μM, scDb-hERG1/β1: 20µg/ml and TTX: 20μM. Corresponding graphs on the bottom panel of actin stress fibres length, filopodia length and cortical actin density. **F)** Staining with anti-cortactin antibody of MCF-7 and SKBR3 cells seeded on FN for 2 hours and treated with E4031: 40μM, scDb-hERG1/β1: 20µg/ml and TTX: 20μM. Corresponding invadopodia graphs on the right panel. At least 20 cells per condition were analysed and all p-values were determined by a Mann–Whitney test for non-parametric values, or for data deviating from normality by a Kolmogorov–Smirnov test. Scale bars: 10 μm. In the graphs, boxes include central 50% of data points, the horizontal lines denote minimum value, median and maximum value. Data are mean values ± s.e.m. (n=3). *P < 0.05, **P < 0.01 and **P < 0.001. Pictures were taken on a confocal microscope (Nikon TE2000, Nikon; Minato, Tokyo, Japan). ImageJ software was used to analyse the images. IF = immunofluorescence.

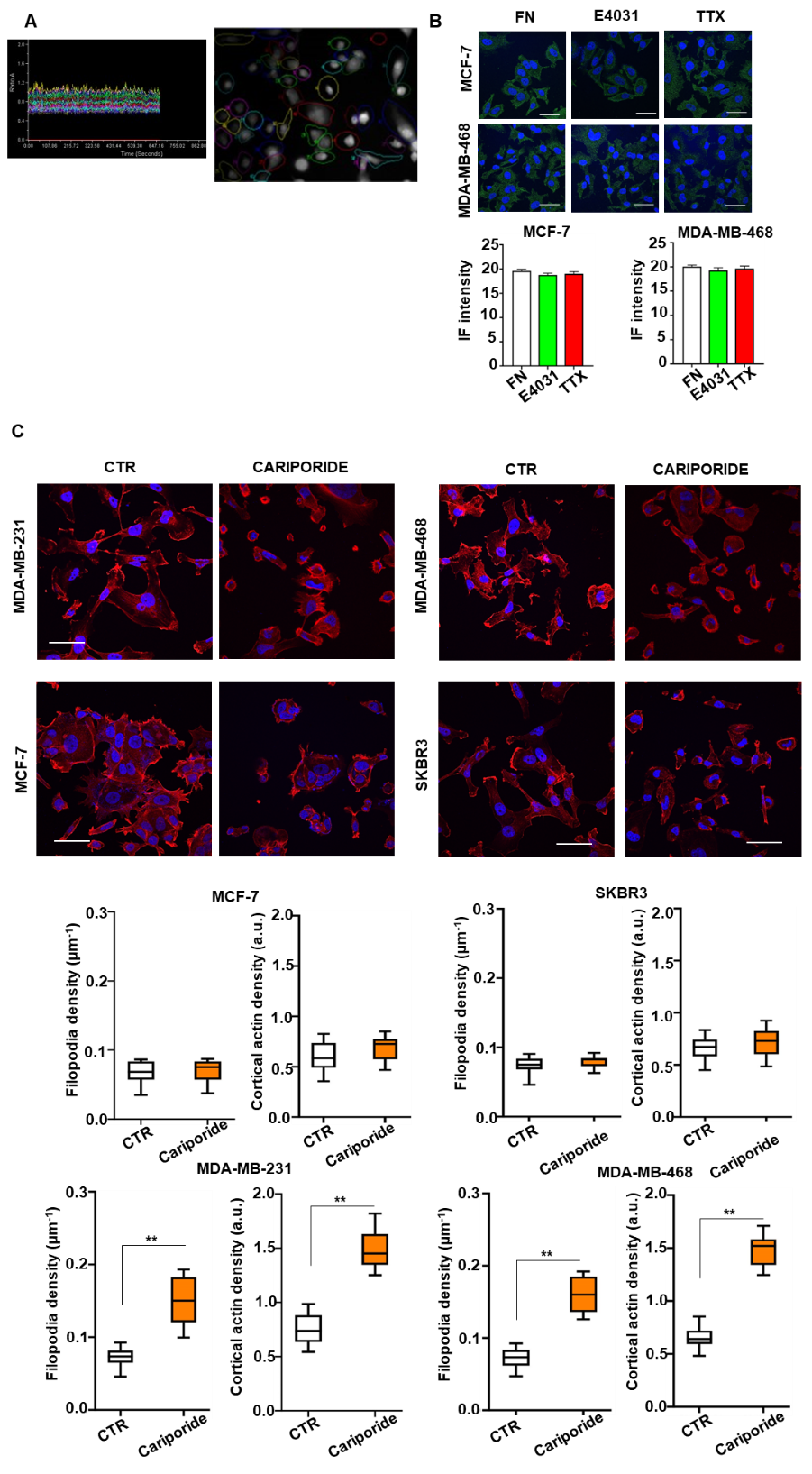

**Supplementary Fig. S8 A)** Measuring of intracellular Ca^2+^ under different conditions. MCF-7 and MDA-MB-468 have been seeded on different type of coating method: FN and BSA for 2h and then intracellular Ca^2+^ was measured as reported in Materials and Methods. Image acquisition (3s frequency) was performed using Metafluor Imaging System (Molecular Devices; Sunnyvale, CA, USA). **B)** pAKT levels were determined through IF. Cells were probed with anti-pAKT antibody followed by secondary antibody conjugated with Alexa Fluor-588 (green). Representative images (scale bar 100 µm) and quantification are reported in the bottom panel. For each condition the fluorescence relative to 20 cells was analysed.

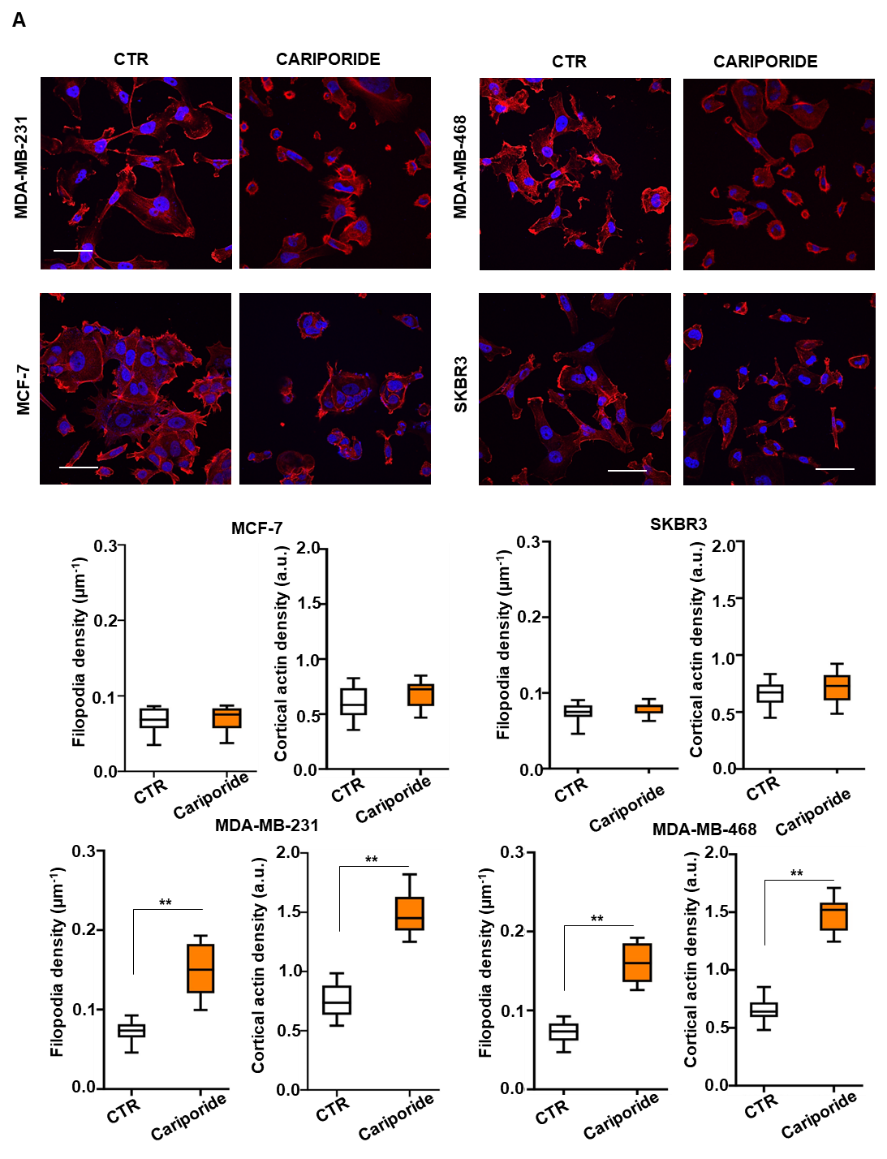

**Supplementary Figure S9. A)** Representative images of phalloidin staining of MCF-7 and SKBR3 cells seeded on FN for 2 hours and treated with Cariporide (5μM). Scale bars: 10 μm. D) Distribution of linear density is calculated as the number of detected filopodia normalize for the cell perimeter. Density distribution of the cortical actin cytoskeleton is also measured. At least 20 cells per condition were analysed. In the graphs, boxes include central 50% of data points, the horizontal lines denote minimum value, median and maximum value. Data are mean values ± s.e.m. (*n*=3). IF = immunofluorescence. FN = fibronectin.

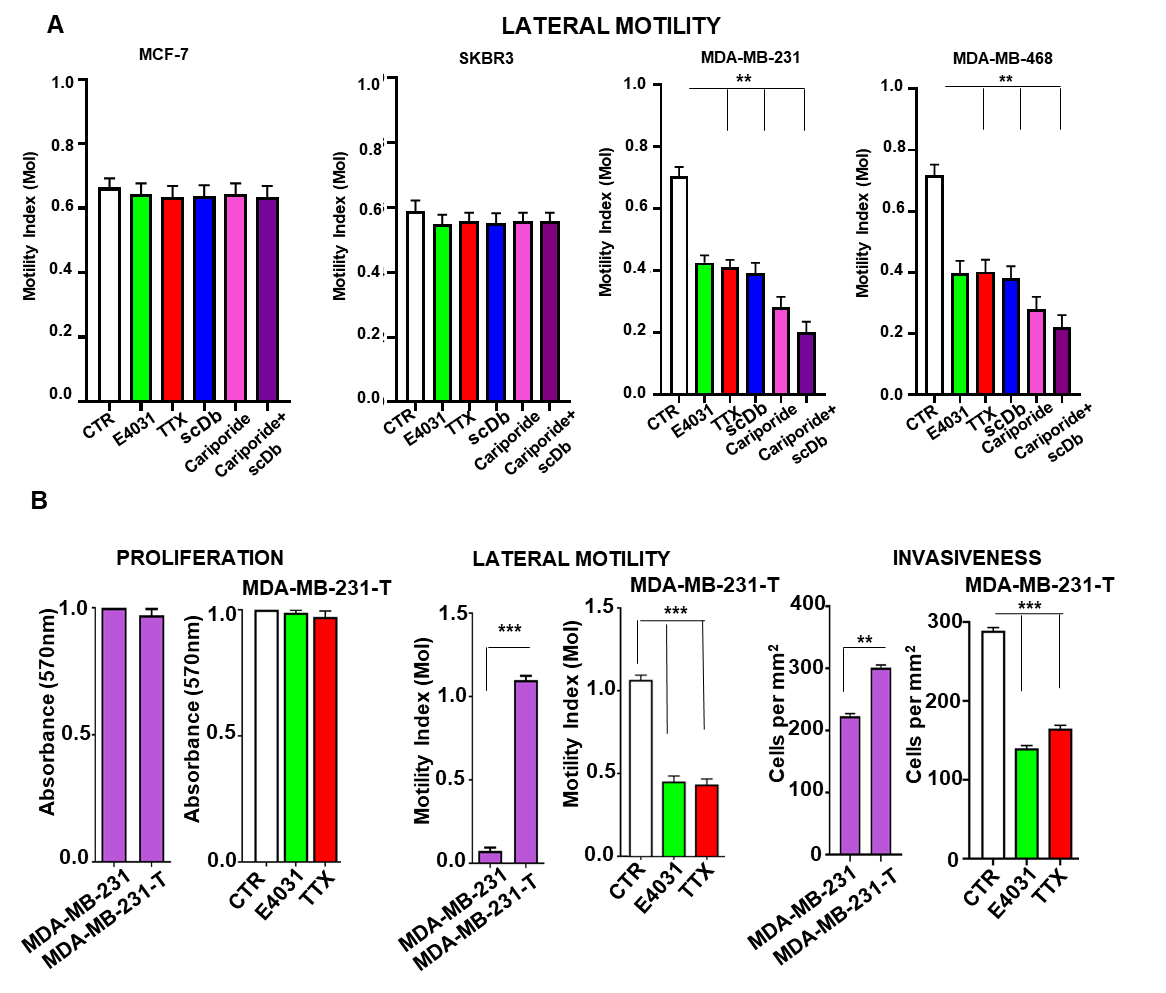

**Supplementary Figure S10. A)** Motility Index (MoI) determined by lateral motility experiments in MCF-7, SKBR3, MDA-MB-231 and MDA-MB-468 cell lines. Cells were treated with E4031 (40μM), tetrodotoxin (TTX) (20μM) and combination E4031 (40μM) + TTX (20μM) for 48 hours. Data are mean values ± s.e.m obtained in three independent experiments. **P < 0.01, ***P < 0.001. **B)** Cell proliferation determined by MTT test in MDA-MB-231 and MDA-MB-231-T cell lines. Cells were treated with E4031 (40μM), tetrodotoxin (TTX) (20μM) for 24 hours and absorbance at 570n was measured. Data are mean values ± s.e.m obtained in three independent experiments. Motility Index (MoI) determined by lateral motility experiments in MDA-MB-231 and MDA-MB-231-T cell lines. Cells were treated with E4031 (40μM), tetrodotoxin (TTX) (20μM) for 24 hours. Data are mean values ± s.e.m obtained in three independent experiments. Invasiveness obtained with matrigel invasion assay in MDA-MB-231 and MDA-MB-231-T. Cells were treated with E4031 (40μM), tetrodotoxin (TTX) (20μM) for 24 hours and cells per mm^2^ were counted. For all experiments, a background level of invasion was calculated, seeding the cells without chemotactic gradient, and a basal level of invasion was then subtracted from the measurements. **P < 0.01 and ***P < 0.001.
